# Structure-Mechanics Principles and Mechanobiology of Fibrocartilage Pericellular Matrix: A Pivotal Role of Type V Collagen

**DOI:** 10.1101/2024.06.26.600498

**Authors:** Chao Wang, Mingyue Fan, Su-Jin Heo, Sheila M. Adams, Thomas Li, Yuchen Liu, Qing Li, Claudia Loebel, Farid Alisafaei, Jason A. Burdick, X. Lucas Lu, David E. Birk, Robert L. Mauck, Lin Han

## Abstract

The pericellular matrix (PCM) is the immediate microniche surrounding resident cells in various tissue types, regulating matrix turnover, cell-matrix cross-talk and disease initiation. This study elucidated the structure-mechanical properties and mechanobiological functions of the PCM in fibrocartilage, a family of connective tissues that sustain complex tensile and compressive loads in vivo. Studying the murine meniscus as the model tissue, we showed that fibrocartilage PCM contains thinner, random collagen fibrillar networks that entrap proteoglycans, a structure distinct from the densely packed, highly aligned collagen fibers in the bulk extracellular matrix (ECM). In comparison to the ECM, the PCM has a lower modulus and greater isotropy, but similar relative viscoelastic properties. In *Col5a1^+/^*^D^ menisci, the reduction of collagen V, a minor collagen localized in the PCM, resulted in aberrant fibril thickening with increased heterogeneity. Consequently, the PCM exhibited a reduced modulus, loss of isotropy and faster viscoelastic relaxation. This disrupted PCM contributes to perturbed mechanotransduction of resident meniscal cells, as illustrated by reduced intracellular calcium signaling, as well as upregulated biosynthesis of lysyl oxidase and tenascin C. When cultured in vitro, *Col5a1^+/^*^D^ meniscal cells synthesized a weakened nascent PCM, which had inferior properties towards protecting resident cells against applied tensile stretch. These findings underscore the PCM as a distinctive microstructure that governs fibrocartilage mechanobiology, and highlight the pivotal role of collagen V in PCM function. Targeting the PCM or its molecular constituents holds promise for enhancing not only meniscus regeneration and osteoarthritis intervention, but also addressing diseases across various fibrocartilaginous tissues.

## INTRODUCTION

In many biological tissues, cells inhabit a structurally distinctive microenvironment, known as the “pericellular matrix” (PCM) or “glycocalyx layer” [1–4]. Owing to its presentation directly at the cell surface, this proximal domain serves as the primary site for the initial assembly of matrix molecules, such as collagen fibrillogenesis and proteoglycan-hyaluronan association [5]. Moreover, the PCM plays a pivotal role in orchestrating cellular activities such as adhesion [6], migration [7,8], growth factor sequestration [9], mechanotransduction [10], stem cell fate [11] and tumorigenesis [12,13]. Notably, the dense glycocalyx layer of PCM has been shown to enhance cancer cell metastasis by facilitating integrin clustering, tension and downstream integrin-dependent signaling pathways [14]. Similarly, in 3D hydrogel cultures, reciprocal interactions between human mesenchymal stem cells (hMSCs) and their PCMs drive stem cell fate and differentiation [15]. In dense connective tissues, the PCM serves as a prevalent structural feature, crucial for transmitting biomechanical cues during their extensive physiological loading. In articular cartilage, the PCM endows a highly negatively charged osmotic environment, essential for chondrocyte homeostasis and responses to compressive and fluid flow mechanical cues [2]. Furthermore, the PCM was identified as the initial point of osteoarthritis (OA) onset, and protecting the PCM could effectively mitigate cartilage degradation [16]. In turn, the clinical significance of cartilage PCM is well appreciated [17,18], as retaining chondrocyte native PCM was shown to improve the quality of regenerative cartilage in OA patients [19]. In bone, the PCM regulates nutrition transport, mechanosensing and survival of osteocytes, thereby modulating the dynamic bone adaptation to mechanical loads [20,21]. In contrast to the cases of articular cartilage and bone, our understanding of the PCM properties and functions in fibrocartilage remains limited.

Fibrocartilage is a family of connective tissues essential for withstanding multiaxial tensile, compressive and shear stresses in vivo. Prominent examples include the knee meniscus, temporomandibular joint (TMJ) condylar cartilage, enthesis, intervertebral disc (IVD) annulus fibrosus and endplate [22]. Reflecting its complex loading conditions, the extracellular matrix (ECM) of fibrocartilage demonstrates a higher degree of compositional and structural heterogeneity compared to both tension-bearing tendon and compression-bearing articular cartilage [23,24]. In fibrocartilage, while the existence of PCM and its potential role in mediating cell-matrix strain transmission has been acknowledged [25–27], understanding of its composition, structure or mechanobiological functions remains elusive. Given the pronounced mechanosensitivity of fibrocartilage cells in homeostasis and pathogenesis [28], targeting the PCM presents a potential avenue for modulating cell mechanotransduction, improving tissue regeneration and mitigating disease progression. Such strategies hold potential for ameliorating diseases associated with fibrocartilage degeneration, such as post-traumatic OA [29], TMJ disorder [30], disc hernia [31] and psoriatic arthritis [32].

This study sought to elucidate the structure-mechanics principles and mechanobiological roles of fibrocartilage PCM. We studied the outer zone of murine meniscus as the model tissue. The meniscus is a loading counterpart of articular cartilage in the knee joint, whose PCM has been studied extensively [33], allowing us to highlight the distinct characteristics of fibrocartilage PCM. Leveraging the murine model enabled us to genetically perturb the PCM constituents, thereby elucidating the molecular activities governing PCM integrity. Specifically, we studied the effects of type V collagen haploinsufficiency using the *Col5a1^+/^*^D^ murine model. Our focus on collagen V stemmed from its pivotal role as the primary nucleation site for initial fibrillogenesis of collagen I [34], the major constituent of meniscus ECM. Moreover, this collagen V-mediated assembly predominantly occurs in the pericellular space [35]. We quantified phenotypic changes in the elastic and viscoelastic nanomechanics of the PCM, along with the resulting alterations in the intracellular calcium signaling activities, [Ca^2+^]*_i_*, in situ, one of the earliest and most fundamental cell responses to biophysical and biochemical stimuli [36]. Next, we elucidated the impact of collagen V reduction on the gene transcriptome of meniscal cells, as well as the integrity and strain-transmission function of newly assembled PCM in vitro. Together, our findings represent the first delineation of the unique molecular and functional properties of the PCM in fibrocartilage, paving the way for enhancing fibrocartilage regeneration through modulation of PCM-mediated matrix assembly and cell mechanotransduction.

## RESULTS

### Distinctive composition, nanostructure and micromechanics of fibrocartilage PCM

In various fibrocartilage tissues, the PCM exhibited distinctive composition and structure compared to the bulk ECM. As shown in bovine and human menisci, as well as human IVD endplate, the PCM was characterized by the preferred distribution of sulfated glycosaminoglycans (sGAGs) and exclusive localization of PCM biomarkers such as perlecan, collagens V and VI (Fig. 1a-c). Also, in these tissues, the PCM comprised randomly oriented, thinner collagen fibrils forming a 3D basket, contrasting with the well-aligned, densely packed collagen fibers in the bulk ECM (Fig. 1a-c). In compression-bearing articular cartilage, although the PCM differed from the bulk ECM by its higher sGAG content, thinner collagen II fibrils and localization of collagen VI, both the PCM and bulk ECM exhibited porous, random collagen fibril networks (Fig. 1d) [37]. This contrasted with fibrocartilage, where these two domains displayed distinct fibrillar organizations (Fig. 1a-c). In tension-bearing tendon, there were no clear marks of distinct PCM domains, and cells were found to reside in spaces between the highly aligned collagen fiber bundles (Fig. 1e). Therefore, the PCM of fibrocartilage was characterized by its unique composition and random, porous collagen fibril architecture, distinguishing it from the bulk fibrous ECM. Similarly, in the outer zone of murine meniscus, the PCM displayed localization of sGAGs and perlecan, as well as preferred distributions of other matrix molecules such as aggrecan, biglycan and collagen VI (Fig. 1f). At the nanoscale, collagen fibrils in the PCM also adopted a porous, randomly oriented basket architecture, contrasting with the circumferentially aligned, densely packed fiber bundles in the bulk ECM (Fig. 1g,h).

**Figure 1.**
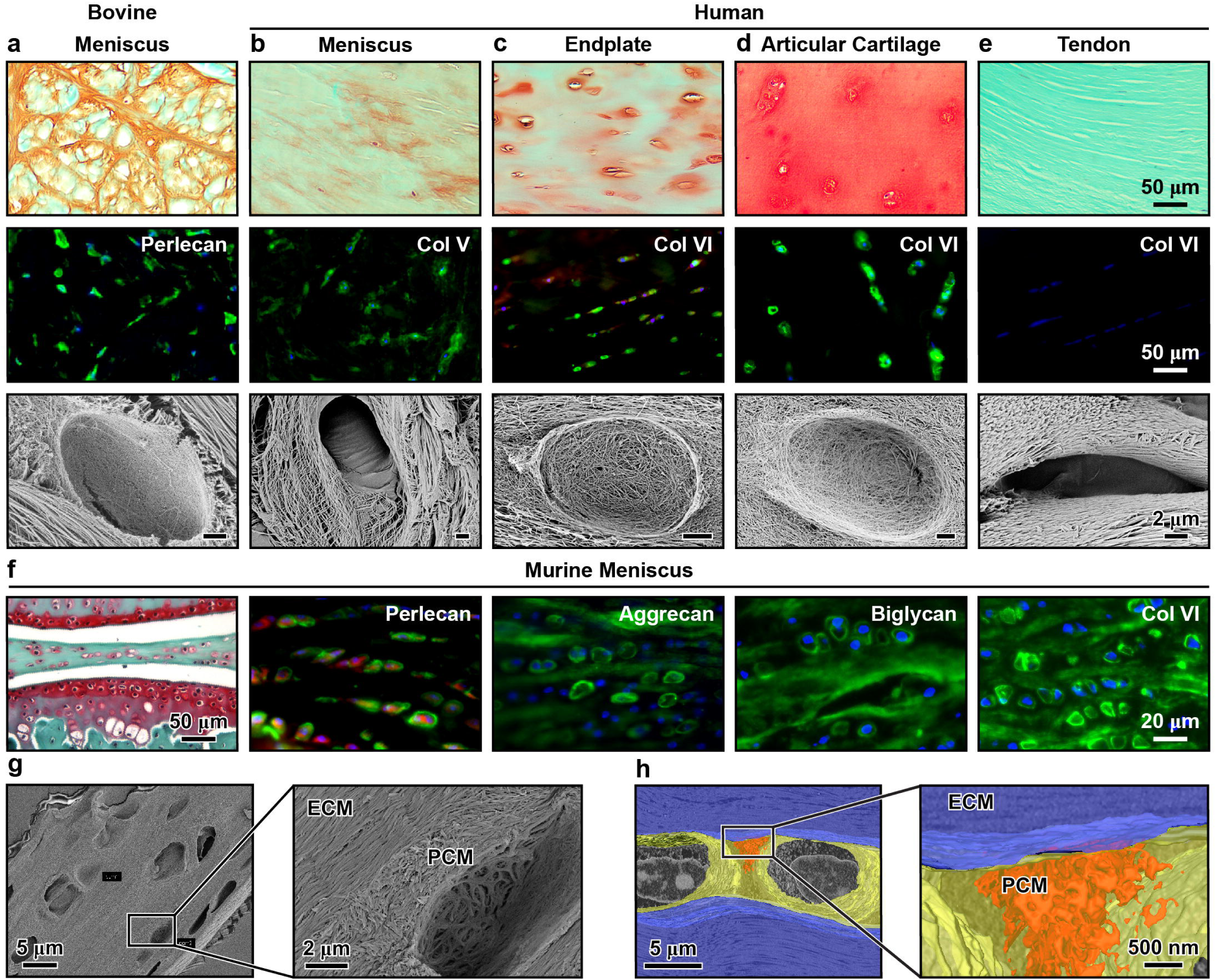
Distinct composition and nanostructure of the fibrocartilage pericellular matrix (PCM) in comparison to the bulk extracellular matrix (ECM). a-e) Molecular distribution and collagen fibril nanostructure of the PCM in fibrocartilage tissues, including a) the bovine meniscus, b) human meniscus, c) human intervertebral disc (IVD) endplate, d) human articular hyaline cartilage, and e) human tendon. Stains include sulfated glycosaminoglycans (sGAGs) via Safranin-O/Fast Green (Saf-O) histology, localization and distribution of matrix molecules in the PCM, such as collagen V (Col V), VI (Col VI) and perlecan (Pln) via immunofluorescence (IF) imaging (blue: DAPI), and collagen fibril network nanostructure visualized by scanning electron microscopy (SEM). SEM results indicate the presence of thinner, more randomly oriented collagen fibrils in the PCM of fibrocartilage, distinct from the highly aligned, thicker collagen fiber bundles in the ECM (*n* = 3 donors, scale bar: 2 µm). f-h) Molecular composition and collagen fibril nanostructure of the meniscus fibrocartilaginous outer zone in 3-month-old mice. f) Localization of sGAGs via Safranin-O histology, perlecan, aggrecan, biglycan and collagen VI via IF imaging (red: cell membrane, blue: DAPI) in the murine meniscus PCM (*n* = 5 mice), g) Porous and randomly oriented collagen fibril network in the PCM, which is distinct from the circumferentially aligned, densely packed collagen fiber bundles in the ECM as measured by e) SEM (*n* = 5) and h) serial block-face (SBF)-3D SEM (orange: higher magnification showing the random fibril architecture in the PCM microdomain, yellow: PCM domain, blue: ECM domain, *n* = 3).

Applying immunofluorescence (IF)-guided AFM nanomechanical tests, we quantified the elastic and viscoelastic properties of the PCM in 3-month-old, adult murine meniscus (Fig. 2a). We measured the effective indentation modulus, *E*_ind_, by fitting each indentation force versus depth (*F-D*) curve to the Hertz model with finite thickness correction [38]. Additionally, we applied the five-element spring-dashpot model to extract the instantaneous modulus, *E*_0_, equilibrium modulus, *E*_∞_, as well as the short-term and long-term relaxation moduli and time constants, (*E*_1_, τ_1_) and (*E*_2_, τ_2_) (τ_1_ < τ_2_), respectively (Fig. 2b). The ratio of equilibrium-to-instantaneous modulus, *E*_∞_/*E*_0_, was used as an indicator of the degree of elasticity. Given the salient structural anisotropy of the fibrous ECM, we tested both horizontal (longitudinal) and vertical (transverse) cryo-sections of the meniscus, for which, the indentation was performed perpendicular and parallel to the ECM fiber axis, respectively (Fig. 2c). Our results highlighted distinct mechanical behaviors of the PCM compared to the fibrous ECM (Fig. 2d-k). First, the PCM exhibited lower *E*_ind_ than the ECM on both horizontal (*E*_ind, PCM_ = 247 ± 31 kPa, *E*_ind, ECM_ = 390 ± 62 kPa, *n* = 9, mean ± 95% CI) and vertical (*E*_ind, PCM_ = 270 ± 71 kPa, *E*_ind, ECM_ = 538 ± 138 kPa, *n* = 7) sections (Fig. 2d). The same contrast was observed for *E*_0_ and *E*_D_ (Fig. 2e,f). Second, in line with its random collagen fibril architecture and high sGAG content (Fig. 1f-h), the PCM displayed isotropic mechanical properties, as we did not note significant orientation-associated differences in either elastic or viscoelastic parameters. This was again different from the pronounced anisotropy observed for the bulk ECM, where indentation parallel to the fiber axis on the vertical section yielded significantly higher *E*_ind_, *E*_D_, *E*_0_, *E*_2_ and longer τ_1_ than that perpendicular to the fiber axis on the horizontal section (Fig. 2c-k). Lastly, despite the pronounced differences in moduli, the PCM exhibited only mild differences in viscoelastic behaviors from the ECM, including a lower *E*_D_/*E*_0_ ratio (Fig. 2i) and a marginally lower τ_1_ (Fig. 2j) on the vertical section, but no other differences.

**Figure 2.**
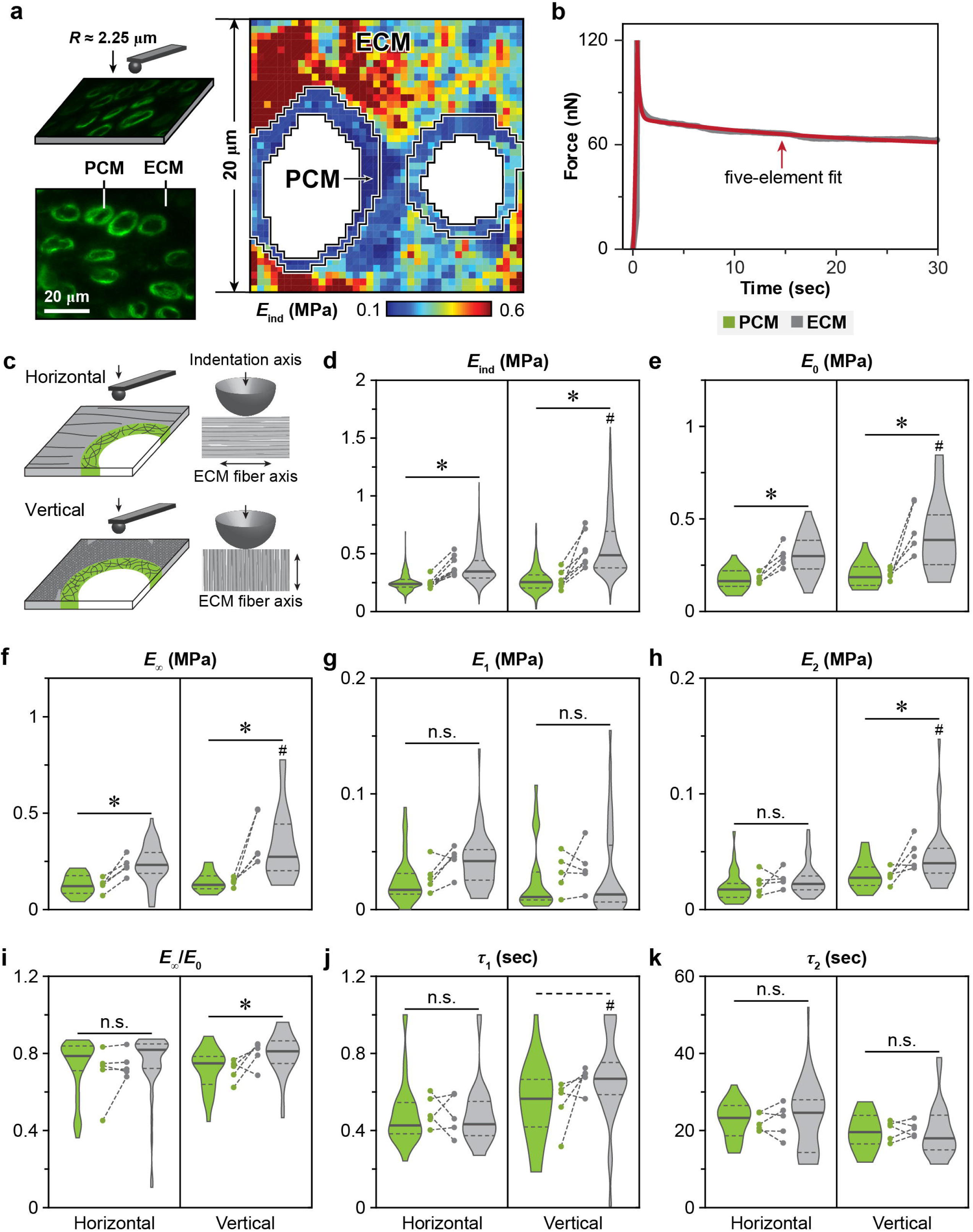
Nanomechanics of the meniscus PCM and ECM via IF-guided AFM-nanomechanical mapping. a) Left panel: Schematic illustration of IF-guided AFM on the horizontal cryo-section of wild-type (WT) murine meniscus using a microspherical tip (*R* ≈ 2.25 µm), where the PCM is immunolabeled with perlecan. Right panel: Representative 20 × 20 µm^2^ indentation micromodulus, *E*_ind_, map on the horizontal cryo-section of untreated murine meniscus. b) Representative 30 sec ramp-and-hold force relaxation curve, measured on the vertical cryo-section of untreated meniscus PCM, and corresponding five-element model fit (R^2^ > 0.99). c) Schematic illustration of the indentation axis with respect to the fiber axis on the horizontal and vertical sections of the meniscus. d-k) Violin plots of elastic and viscoelastic micromechanical properties of 3-month-old WT murine meniscus PCM and ECM on horizontal and vertical cryo-sections, including d) indentation micromodulus, *E*_ind_, e) instantaneous modulus, *E*_0_, f) equilibrium modulus, *E*_∞_, g) short-term viscoelastic relaxation modulus, *E*_1_, h) long-term viscoelastic relaxation modulus, *E*_2_, i) elasticity ratio, *E*_D_/*E*_0_, j) short-term relaxation time constant, τ_1_, and k) long-term relaxation time constant, τ_2_. Data are pooled from ≥ 580 positions tested for *E*_ind_, ≥ 25 positions for viscoelastic properties for each group (*n* = 5 mice). Each circle represents the average value measured from one animal. *: *p* < 0.05, dashed line: *p* < 0.10, n.s.: not significant between PCM and ECM under the same indentation orientation, ^#^: *p* < 0.05 between the horizontal and vertical sections for each of the PCM or ECM domain.

### Collagen V regulates the fibril nanostructure and micromechanics of meniscus PCM

Next, using the *Col5a1^+/−^* model, we queried how perturbations in PCM molecular constituents impacts its nanostructure and biomechanical properties. In both immature (3-week-old) and adult (3-month-old) *Col5a1^+/−^* murine menisci, we found preferred localization of collagen V in the PCM, alongside a noticeable reduction of collagen V compared to their age-matched WT controls (Fig. 3a). Despite this reduction, there were no marked alterations in the preferred distribution of large proteoglycan, aggrecan (Fig. 3b) or sGAGs (Fig. 3c) within the PCM. Applying TEM to 3-month-old menisci, we found disrupted fibril nanostructure in the *Col5a1^+/−^* meniscus PCM, marked by substantially thickened fibrils with higher heterogeneity (variance) (Fig. 3d-f). Also, in both genotypes, fibrils in the PCM were thinner than those in the bulk ECM. Thus, with the reduction of collagen V, the PCM developed impaired fibrillar architecture, but retained its compositional and structural distinction from the bulk ECM.

**Figure 3.**
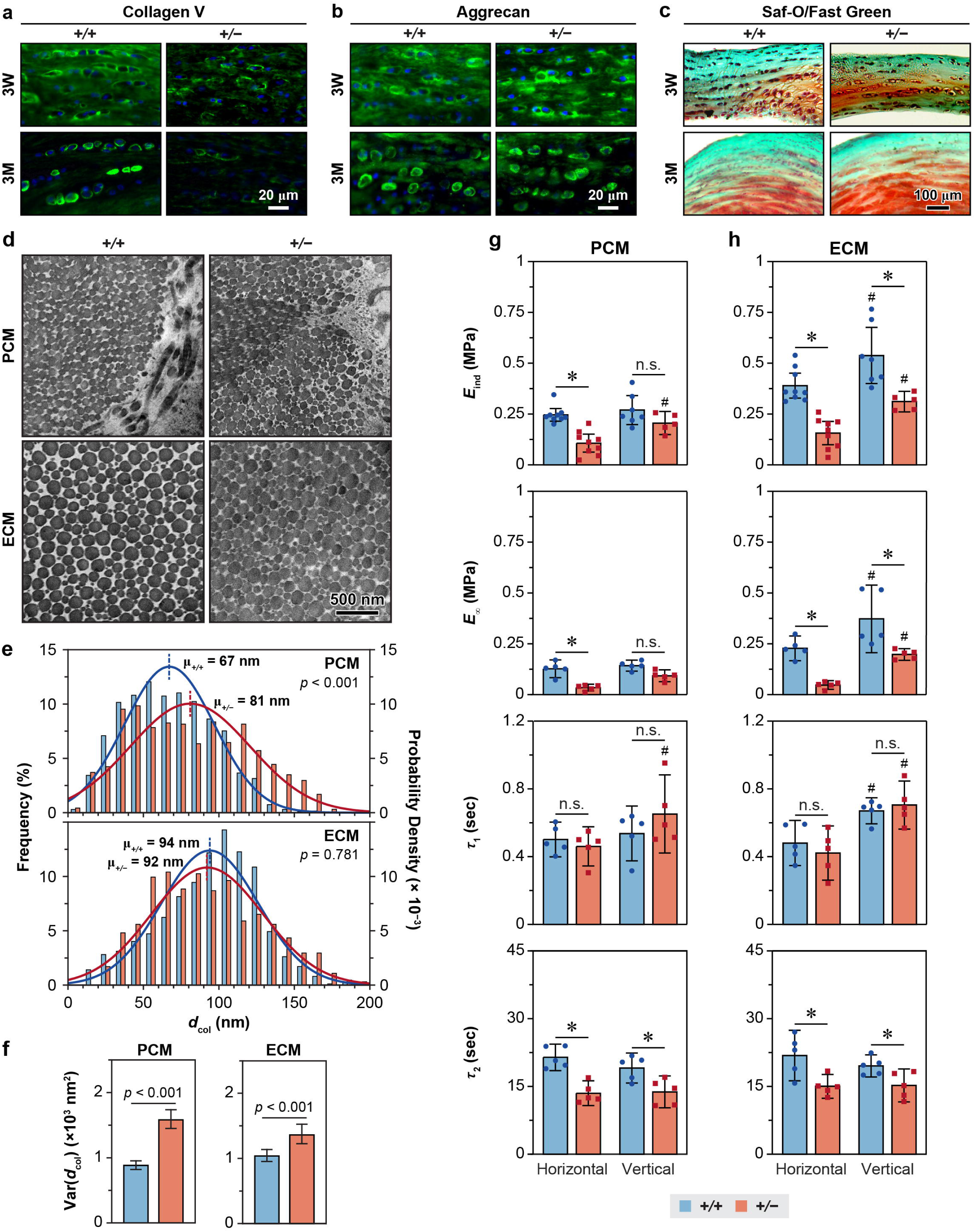
Impact of collagen V haploinsufficiency on the nanostructure and micromechanics of the meniscus PCM. a) Safranin-O/Fast Green histology illustrates no appreciable difference in the morphology and sGAG staining between wild-type (+/+) and *Col5a1^+/−^* (*+/−*) murine meniscus at 3 weeks and 3 months of ages (*n* ≥ 6 mice). b,c) Immunofluorescence (IF) images show b) reduced collagen V content and c) no apparent changes of aggrecan content in the meniscus of *Col5a1^+/−^*relative to that of the WT control at 3 weeks and 3 months of ages (*n* ≥ 6). d) Representative TEM images of collagen fibrils in the PCM and ECM of WT and *Col5a1^+/−^* meniscus, as imaged on tissue vertical sections. e) Histogram of fibril diameter distribution (≥ 640 fibrils from *n* = 5 mice for each genotype within each region). Shown together are the normal distribution, *N*(*µ*, σ^2^), fits to the fibril diameter distributions (for each fit, values of *µ* and σ correspond to the mean and standard deviation of fibril diameters). f) Comparison of fibril heterogeneity (variance) in the PCM and ECM of WT and *Col5a1^+/−^*meniscus (mean ± 95% CI, ≥ 640 fibrils from *n* = 5 mice). g,h) Elastic and viscoelastic micromechanical properties of the murine meniscus for g) PCM and h) ECM on horizontal and vertical cryo-sections, for WT and *Col5a1^+/−^* groups, including the indentation micromodulus, *E*_ind_, equilibrium modulus, *E*_∞_, short term, τ_1_, and long term, τ_2_, relaxation time constants (mean ± 95% CI, *n* ≥ 5). Each data point represents the average value measured from one animal. *: *p* < 0.05 between WT and *Col5a1^+/−^* groups under same orientation and region, ^#^: *p* < 0.05 between the horizontal and vertical sections for same treatment and region. Results from panels d-h were measured from murine menisci at 3 months of age.

In line with the disrupted fibril nanostructure, the reduction of collagen V resulted in altered PCM micromechanics, including decreased *E*_ind_, *E*_0_ and *E*_D_ for horizontal section, as well as faster τ_2_ for both sections. Notably, the PCM of *Col5a1^+/−^*menisci exhibited salient anisotropy in *E*_ind_, *E*_0_, *E*_2_ and τ_1_ (Fig. 3g and Fig. S3), contrasting with the isotropic micromechanical nature of the WT meniscus PCM. Meanwhile, besides its impact on the PCM, reduction of collagen V also influenced the structure and biomechanics of the bulk ECM. The ECM of *Col5a1^+/−^*menisci exhibited higher fibril diameter variance despite having similar average diameters compared to the WT ECM (Fig. 3f). Additionally, the *Col5a1^+/−^* ECM showed decreased *E*_ind,_ *E*_0_ and *E*_∞_ and faster τ_2_ (Fig. 3h), indicating a role of collagen V in overt tissue integrity beyond its impact on the PCM.

### Reduction of collagen V disrupts PCM-mediated meniscal cell mechanotransduction

We then tested if disruption of PCM integrity due to collagen V reduction perturbs the mechanotransduction of resident meniscal cells. We studied the intracellular calcium signaling activities, [Ca^2+^]*_i_*, of freshly dissected medial meniscus explants in situ. Since the size and irregular shape of the meniscus limits the application of well-defined mechanical strains, we tested the responses in both physiological (isotonic) and osmotically instigated (hypo- and hypertonic) Dulbecco’s Modified Eagles Medium (DMEM). Using this setup, we examined both WT and *Col5a1^+/−^*menisci at 3 weeks and 3 months of age. For all tested groups, we observed spontaneous [Ca^2+^]*_i_* oscillations, from which we extracted temporal [Ca^2+^]*_i_* parameters, including the percentage of responding cells, %*R*_cell_, the total number of [Ca^2+^]*_i_* peaks, *n*_peak_ from each responding cell over a 15-minute observation period (Fig. 4a). Compared to the WT, *Col5a1^+/−^* cells exhibited substantially reduced [Ca^2+^]*_i_* activities, with lower %*R*_cell_ at all osmolarities and both ages, except for the hypertonic condition at 3 weeks (Fig. 4b). Additionally, *Col5a1^+/−^* meniscal cells displayed lower *n*_peak_ than the WT in isotonic DMEM at 3 weeks, and hypotonic DMEM at 3 months (Fig. 4c). For both genotypes, immature cells at 3 weeks showed more active [Ca^2+^]*_i_*responses than adult cells at 3 months, suggesting that collagen V deficiency did not abrogate maturation-associated changes in cell mechanosensing. Furthermore, similar to WT cells, *Col5a1^+/−^* cells demonstrated salient osmolarity-dependence, evidencing loss of collagen V did not abolish cell mechanosensing in response to changes in their osmotic environment.

**Figure 4.**
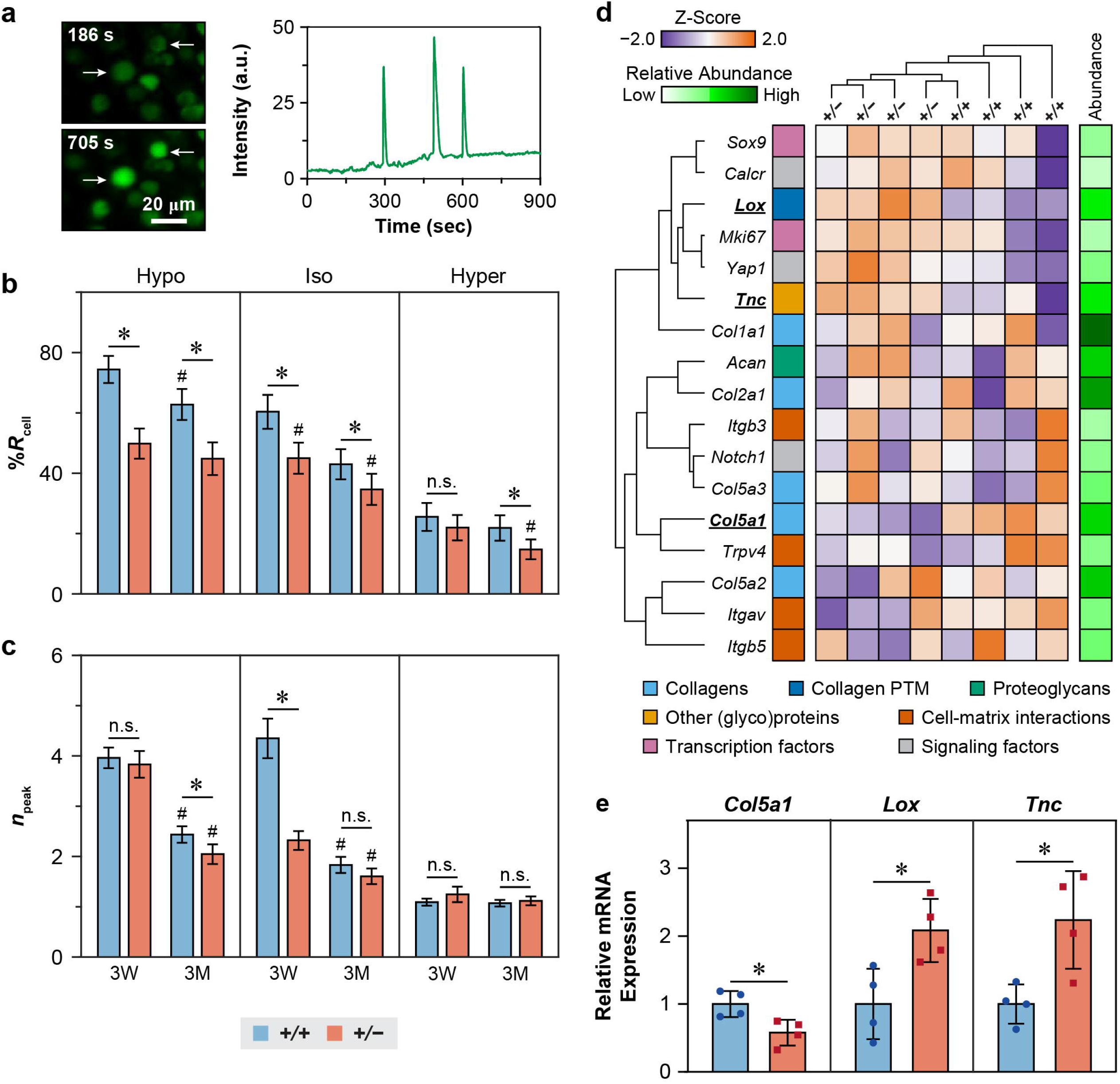
Impact of collagen V haploinsufficiency on the in situ mechanotransduction and gene expression of meniscus cells. a) Left panel: Representative confocal images of [Ca^2+^]*_i_* signaling of the untreated meniscus explant in isotonic DMEM. Right panel: Representative [Ca^2+^]*_i_*oscillation fluorescence intensity curve over a 15-min time frame of a single cell from untreated murine meniscus in hypotonic DMEM. b,c) Comparison of in situ [Ca^2+^]*_i_* signaling parameters between wild-type (WT, +/+) and *Col5a1^+/−^* (*+/−*) at 3 weeks and 3 months of ages in hypotonic, isotonic and hypertonic DMEMs at 37°C, including b) the percentage of responding cells, %*R*_cell,_ and c) number of peaks within 15-min testing time frame, *n*_peak_ (mean ± 95 % CI, ≥ 65 responsive cells pooled from *n* = 4 mice for each group. *: *p* < 0.05 between genotypes under same age and osmolarity, n.s.: not significant. ^#^: *p* < 0.05 between ages within same genotype and osmolarity). d) Heatmap of unbiased clustering of multiplexed selected genes for +/+ and *+/−* menisci at 3 weeks of age (*n* = 4 for each genotype). Color legend for relative mRNA expression. For the heatmap with a full gene list used in the cluster analysis, see Figure S4. e) Comparison of the expression of selected genes between 3-week-old WT and *Col5a1^+/−^* meniscus measured by quantitative PCR (mean ± S.D., *n* = 4, *: *p* < 0.05). Each data point represents the average value measured from one biological replicate of cells pooled from 2 animals.

Considering that meniscal cells were more mechanoresponsive at the immature age of 3 weeks (Fig. 4b,c), we tested how the disrupted PCM alters the expressions of key mechanotransduction genes by applying NanoString multiplex gene expression analysis to 3-week-old meniscus (*n* = 4 biological replicates/genotype), which directly counts RNA molecules within a custom-built gene panel set [39]. A panel set of 103 genes were tested, including major genes of collagens, collagen post-translational modification (PTM), proteoglycans and other matrix proteins/glycoproteins, cell-matrix interactions, surface markers, transcriptional factors, as well as biomarkers for major mechanosensitive pathways (Table S5). Besides the expected decrease in *Col5a1*, we also found significant increases in *Lox* and *Tnc* (Fig. 4d and Fig. S4), which encode lysyl oxidase (LOX) and tenascin-C, respectively. We then confirmed these impacted genes by qPCR (Fig. 4e). However, we did not find significant changes in all other tested genes. Together, these results supported that the altered PCM microenvironment resulted in perturbed mechanosensing of the resident meniscal cells, which could, in turn, alter the expressions of other matrix molecules.

### Reduction of collagen V impairs the assembly and function of meniscus neo-PCM

Finally, we investigated whether collagen V regulates the PCM assembly and cell-PCM interactions at the cellular level. We extracted meniscal cells from 3-week-old WT and *Col5a1^+/−^* menisci, and cultured them in vitro. Applying metabolic azidohomoalanine (AHA) “click-labeling” [40], we assessed the distribution of newly synthesized proteins within the nascent matrix, or “neo-PCM”, surrounding individual meniscal cells for up to 7 days in 2D culture. Comparing the two genotypes, we observed no significant differences in the thickness of neo-PCM, except for a mildly higher thickness for *Col5a1^+/−^*cells at day 3 (Fig. 5a,b). This suggests that reduction of collagen V did not substantially alter the distribution of newly synthesized proteins during neo-PCM formation; however, the method is unable to discern the amount of new proteins in the neo-PCM. On the other hand, *Col5a1^+/−^*meniscal cells synthesized a higher amount of LOX and tenascin-C proteins (Fig. 5c), consistent with their increased gene expression found in vivo (Fig. 4d,e). We then applied AFM-nanoindentation to individual cells with their associated neo-PCM. At day 0, with the absence of neo-PCM, WT and *Col5a1^+/−^* cells showed similar apparent modulus (Fig. 5d). By day 7, there was an increase in apparent modulus for both genotypes, likely evidence of the deposition of neo-PCM surrounding the cells. However, the micromodulus was lower for collagen V-deficient neo-PCM at day 7 (Fig. 5d), despite similar thicknesses (Fig. 5b) and higher LOX content (Fig. 5c). This lower modulus also aligns with the lower modulus found in the native PCM of *Col5a1^+/−^* meniscus (Fig. 3g), affirming the crucial role of collagen V in mediating the assembly and quality of meniscus PCM.

**Figure 5.**
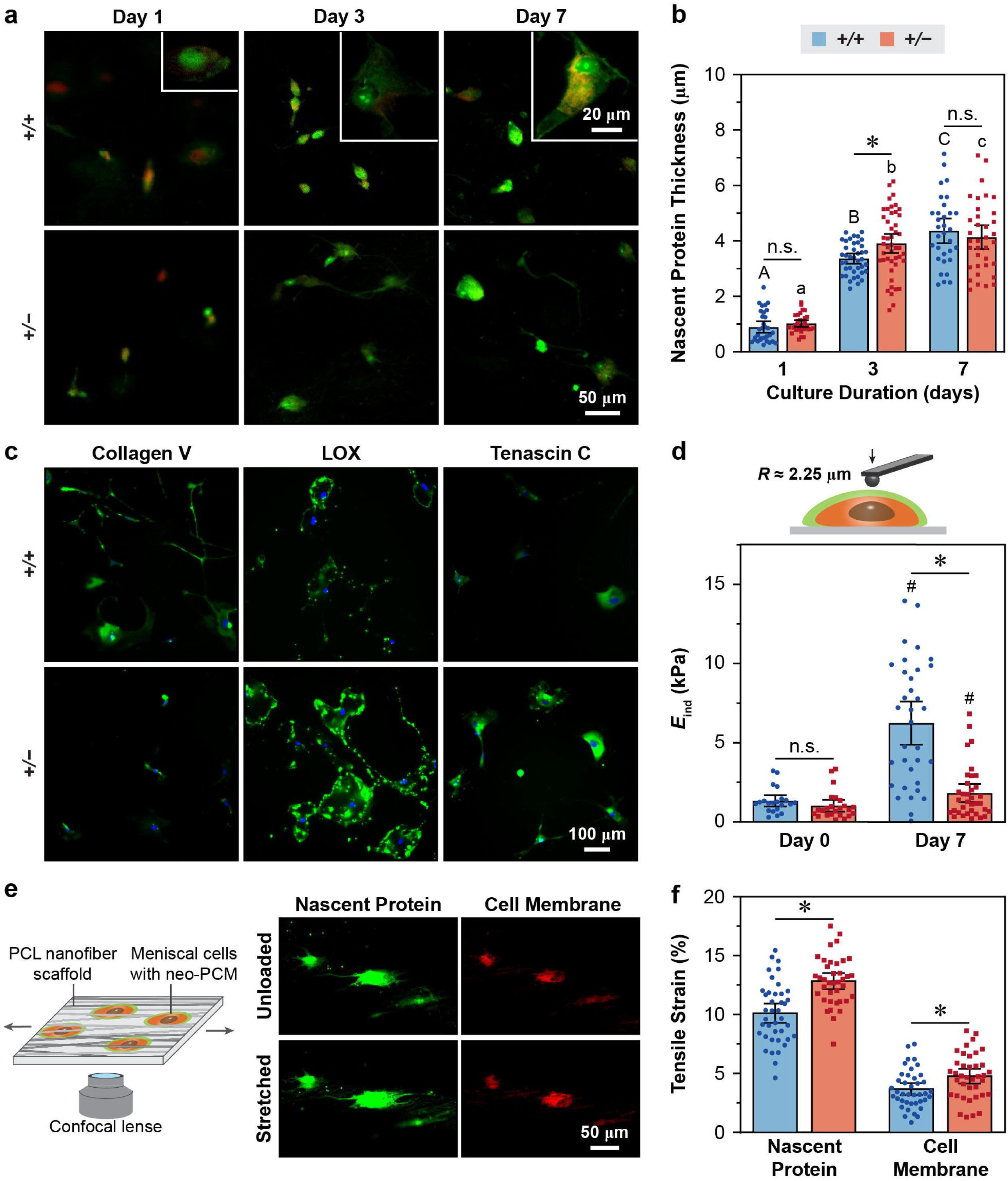
Impact of collagen V haploinsufficiency on the assembly and strain transmission function of neo-pericellular matrix (neo-PCM) synthesized by meniscus fibrochondrocytes. a) Representative fluorescence images of nascent proteins (green) deposited by wild-type (WT, +/+) and *Col5a1^+/−^*(*+/−*) meniscal cells cultured for 7 days, as measured by metabolic click-labeling of azidohomoalanine (AHA). b) Quantification of the accumulated nascent protein thickness deposited by WT and *Col5a1^+/−^*meniscus cells cultured for 1, 3 and 7 days, respectively (mean ± 95 % CI, pooled from *n* ≥ 30 cells extracted from the menisci of ≥ 3 mice from 3 independent experiments, *: *p* < 0.05 between genotypes, n.s.: not significant, different letters indicate significant differences between culture duration within each genotype). c) Immunofluorescence (IF) staining images of collagen V, lysyl oxidase (LOX) and tenascin-C deposited by WT and *Col5a1^+/−^* meniscus cells cultured for 7 days. d) Effective AFM-nanoindentation micromodulus, *E*_ind_, of the neo-PCM with cells shows reduced modulus for the *Col5a1^+/−^*group at 7 days, but not 0 days, of culture (mean ± 95% CI, *n* ≥ 20 cells extract from the menisci of ≥ 3 mice, *: *p* < 0.05 between genotypes for each culture time point, ^#^: *p* < 0.05 between culture time points within each genotype, n.s.: not significant). e) Left panel: Schematic illustration of confocal imaging on the tensile stretch of meniscal cells with neo-PCM embedded in a PCL nanofiber scaffold. Right panel: Representative confocal images of nascent protein (green) and cell membrane (red) of individual meniscus fibrochondrocytes under 0% and 20% applied tensile stretch to the underlying PCL scaffold after culture for 7 days. f) Comparison of strain transfer from 20% applied tensile stretch to nascent protein and cell membrane between +/+ and *+/−* meniscal cells after 7-day cultured within the PCL scaffold (mean ± 95 % CI, *n* ≥ 38 cells extracted from menisci of ≥ 3 mice, *: *p* < 0.05 between genotypes). Panels a-f were obtained from meniscal cells extracted from 3-week-old mice.

To test if this weakened neo-PCM alters strain transmission, we cultured meniscal cells with an aligned porous poly(ε-caprolactone) (PCL) nanofiber scaffold for 7 days to enable the development of neo-PCM, and tracked the neo-PCM formation via AHA click-labeling. By day 7, cells formed a composite with the neo-PCM (Fig. 5e), which were embedded in the aligned scaffold, mimicking the structure of native meniscus with cell-PCM entrapped within a fibrous ECM (e.g., Fig. 1). Under 20% applied tensile strain to the PCL scaffold, the tensile strain along the stretch direction for *Col5a1^+/−^* neo-PCM was significantly higher than that of WT (Fig. 5f). Similarly, *Col5a1^+/−^* cells also showed higher strain than the WT. These results suggest that the collagen V-deficient neo-PCM exhibited impaired protection of resident cells under external tensile stretch, corroborating its reduced micromodulus (Fig. 5d).

## DISCUSSION

This study underscores the PCM as a structurally distinctive microdomain across multiple fibrocartilaginous tissues (Fig. 1). Comprising a porous, random fibrillar architecture in 3D with thinner collagen fibrils with preferred localization of proteoglycans and their sGAGs, the PCM stands in stark contrast to the densely packed, aligned collagen fibers of the ECM (Fig. 1). Studying murine meniscus, we highlight the distinctive structure-mechanics relationship of fibrocartilage PCM (Fig. 6). First, the lower modulus of the PCM relative to the ECM (Fig. 2d) can be attributed to its less organized fibrillar structure, such as thinner fibrils and lower density. While the higher sGAG content in the PCM provides additional compressive resistance through fixed charge-endowed osmotic swelling and its confining effects of limiting collagen fibril bending or buckling [41], the impact of less organized fibrillar structure seems to outweigh this contribution, resulting in a net lower modulus. Second, the isotropic mechanical properties of PCM stand in contrast to the salient anisotropy of the ECM (Fig. 2d). For the PCM, spherical nanoindentation results in the same deformation mode on both horizontal (longitudinal) and vertical (transverse) sections, involving mutual engagement of the collagen fibril network and sGAGs [41]. This isotropy arises from the random fibrillar architecture as well as the presence of sGAGs, which not only provide isotropic osmotic swelling pressure, but stretches collagen fibrils in an isotropic manner [42]. In contrast, for the fibrous ECM, based on previous studies on tendon, cervix and bovine meniscus [33,43], the anisotropy originates from different deformation modes of collagen fibers under spherical nanoindentation. Indentation parallel to the fiber axis (on horizontal section) leads to mutual engagement of multiple fibrils undergoing laterally confined multiaxial compressive response, similar to a material continuum. Indentation normal to the fiber axis (on vertical section) results in local fibril uncrimping and bending, which involve smaller shear strain transfer amongst the fibrils and are more compliant.

**Figure 6.**
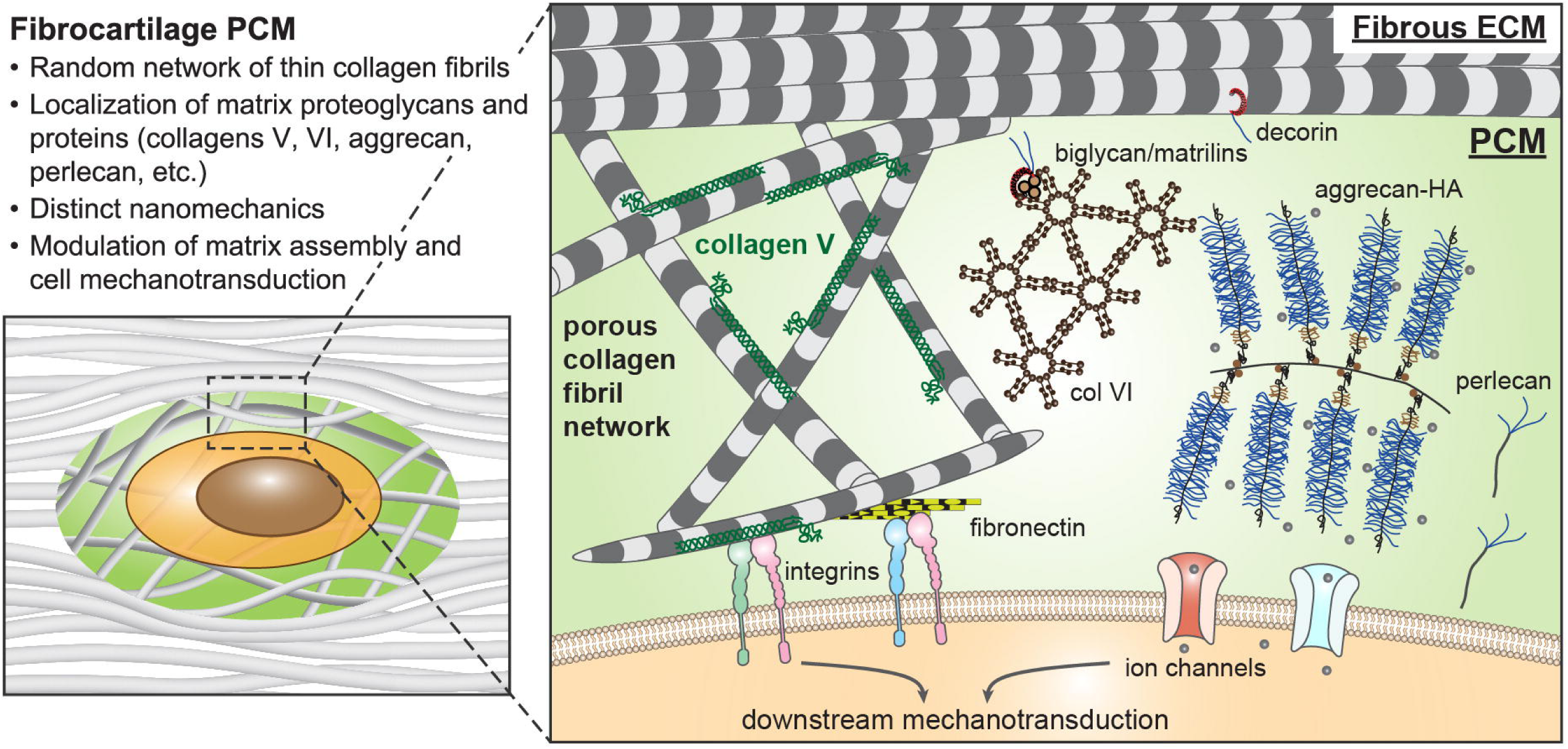
Schematic illustration of the structure and composition of fibrocartilage PCM. In fibrocartilage such as the meniscus, the PCM is characterized by a random network of thinner collagen fibrils and localization of regulatory matrix proteins and proteoglycans, resulting in isotropic nanomechanical properties that is different from the bulk fibrous ECM. Within the PCM, the fixed negative charges endowed by proteoglycans such as aggrecan directly contribute to its nanomechanical properties. In addition, collagen V is highly concentrated in the PCM, and plays pivotal roles in the fibril assembly and mechanobiological functions of the PCM.

Despite their marked differences in structure and modulus, the PCM and ECM exhibit similar relative time-dependent mechanical characteristics and time constants (Fig. 2). The only notable difference is a mildly higher elasticity ratio *E*_D_/*E*_0_ of the ECM on the vertical section (Fig. 2i), likely due to its higher fiber packing density. In soft biological tissues, time-dependent mechanics is governed by two mechanisms. Fluid flow-induced poroelasticity dissipates energy through water-solid matrix interactions [44], while intrinsic viscoelasticity dissipates energy through molecular frictions during matrix macromolecular re-configuration and is independent of fluid flow [45]. Under spherical nanoindentation with an indenter tip of *R* ≈ 2.25 µm and a maximum indentation depth of *d*_max_ ≈ 100-200 nm, the characteristic fluid flow length, *L*_P_ ≈ *R* × arccos[(*R* – *d*_max_)/*R*] = 0.6-1.0 µm. For the meniscus matrix, the hydraulic permeability, *k* is ∼ 1 × 10^-15^ *m*^4^/(*N*·*s*) [46], and our nanoindentation results suggest an aggregated modulus, *H*_A_ ∼ 0.1-0.2 MPa (Fig. 2f), the poroelastic time constant τ_P_ is ∼ *L*_P_^2^/(*H*_A_*k*) ≈ 2-10 msec << 0.1 sec [47]. Therefore, at this length scale, poroelasticity is expected to contribute minimally to the observed relaxation behaviors, which are dominated by intrinsic viscoelasticity. In the PCM, although thinner fibrils may expect higher molecular mobility and faster relaxation than the densely packed ECM fibers, this effect is likely offset by the confining effects of sGAGs that slow collagen fibril movement [41]. These similarities in the relative viscoelasticity across different tissue compartments has been hypothesized to minimize local stress and strain redistribution, and reduce stress concentration at their interfaces during time-dependent deformation [33]. Such self-consistency in time-dependent mechanics has been observed in various heterogeneous tissue structures, including bovine meniscus across different structural units at the microscale [33] and across various anatomical sites at the mesoscale [48], as well as bovine cartilage throughout tissue depth [49].

The distinct structure-mechanics principles of the meniscus PCM represent specialized mechanical adaptation of fibrocartilage to its complex loading environment. In articular cartilage, the PCM adapts a similar structure of porous collagen II fibril networks and proteoglycans as the ECM, and its higher sGAG content compared to the ECM is central to chondrocyte mechanosensing of compressive loads [50,51] and protection against overloading. In tendons and ligaments, cells reside within the fibrous matrix without an intermediary PCM, except in specific case such as wrap-around tendon that also sustains compression [52] and injured tendon undergoing aberrant remodeling [53]. Fibrocartilage, including the meniscus, sustains a combination of tensile and compressive loads. In the knee joint, the meniscus sustains tensile hoop stress to provide joint stability [54] while transmitting femur-tibia compressive stress to reduce cartilage loading [55]. Under physiological joint loading, stretching of the circumferential ECM fibers results in multiaxial stress to the cell, tension along the fiber axis, and transverse compression normal to the fiber axis [56]. Therefore, sGAGs in the PCM can resist transverse compression [57], and the randomly oriented collagen fibril network can redistribute tensile stress to reduce force transmitted to the cell [58]. Both constituents thus contribute to attenuating cell strain and protect cells from tensile and compressive overloading (Fig. 6). Indeed, our findings demonstrate that perturbing the neo-PCM assembly by reducing a critical PCM constituent, collagen V, leads to higher deformation of meniscal cells under applied tension (Fig. 5d-f), indicating impaired protection to resident cells.

We show collagen V as a critical constituent maintaining the meniscus PCM integrity, both in native tissue (Fig. 3) and during in vitro cell culture (Fig. 5). In *Col5a1^+/−^* meniscus PCM, the observed thickening and increased heterogeneity of collagen fibrils (Fig. 3d-f) can be attributed to the canonical role of collagen V in mediating collagen I fibrillogenesis in the pericellular space [34,59]. This altered fibril structure could also disrupt the integration between the fibrillar network and proteoglycans, leading to impaired strain transfer between the fibrils and reduced osmotic pressurization, ultimately resulting in lower modulus and loss of mechanical isotropy (Fig. 3g). This disrupted fibril assembly and integration with proteoglycans also contribute to higher molecular mobility for fibril re-arrangement under stress, resulting in faster relaxation time, τ_2_ (Fig. 3g). Additionally, reduction of collagen V alters the structure and mechanics of the fibrous ECM, despite its lower concentration therein (Fig. 3h). This effect could also arise from the pivotal role of collagen V mediating PCM fibril assembly. During tissue development, early collagen fibrils are deposited and assembled in the pericellular space close to the cell body [60]. These fibrils provide the template for growth into the mature fibrous ECM that is further away from the cells [61]. The pre-deposited defects due to collagen V deficiency in the pericellular fibrils thus lead to compromised ECM formation. Therefore, collagen V not only directly regulates the PCM fibril assembly, but also mediates the overall integrity of the meniscus matrix. While the *Col5a1^+/−^*mice can serve as the model for disrupted meniscus PCM, this constitutive heterozygous model does not allow us to query the impact of complete collagen V ablation due to embryo lethality of homozygous *Col5a1^-/-^* mice [62]. Our future work will establish targeted collagen V knockout models by cross-breeding *Col5a1^f/f^* mice [63] with conditional inducible Cre models (e.g., *Col1a2Cre^ER^*) [64] to control the timing of *Col5a1* ablation in the meniscus. This will enable us to achieve near-complete ablation of collagen V without lethality and to delineate its activities in PCM versus ECM during different stages of development and remodeling. Despite this limitation, given the immediate PCM-cell contact, the observed changes in cell-matrix mechanosensing in *Col5a1^+/−^* meniscus (Fig. 4) are expected to be manifested through the effects of collagen V on the PCM, rather than the ECM that is farther removed from cells.

We further illustrate the pivotal role of collagen V in PCM-mediated cell mechanosensing. Specifically, the demoted in situ [Ca^2+^]*_i_*activities observed in *Col5a1^+/−^* meniscal cells (Fig. 4b,c) underscore the impaired mechanosensing of the disrupted PCM. In contrast, multiplex gene expression analysis does not yield profoundly altered downstream signaling pathways (Fig. 4d). Cellular mechanotransduction involves a cascade of molecular events, in which, cells sense and respond to mechanical stimuli by the rearrangement of cytoskeleton, and cytoskeletal forces are transmitted into the nucleus. Consequently, nuclear forces lead to chromatin reorganization and alterations in the epigenetic landscape, thereby regulating gene expression [65,66]. Our results suggest that meniscal cells possess an inherently robust mechanosensing framework for maintaining normal homeostasis resilient to alterations in PCM properties and transmembrane [Ca^2+^]*_i_* activities. This may represent an adaptive biological process for meniscal cells to maintain normal homeostasis under extensive physiological loadings. For example, following sciatic nerve resection surgery, murine meniscus was found to undergo normal postnatal growth despite substantial disruptions in joint loading and gaits [67]. Also, in other tissues, collagen V plays direct biological roles such as maintaining the muscle satellite stem cell niche through the Notch-Col V-Calcr axis [68], influencing fibroblast growth factor-2 (FGF-2)-mediated angiogenesis [69], as well as mediating cardiac and dermal wound healing by impacting integrin expression [70–72] and transforming growth factor-β (TGF-β) signaling in fibroblasts [73]. However, our multiplex gene expression analysis of the *Col5a1^+/−^* meniscus shows no substantial changes in these pathways (Fig. S5), suggesting either collagen V does not perform such roles in the meniscus, or that its biological activities are retained despite heterozygous reduction. Nevertheless, it is noteworthy that this impaired PCM still results in the up-regulation of two crucial matrix proteins, LOX and tenascin-C, in both native tissue (Fig. 4d,e) and in vitro cell culture (Fig. 5c). LOX is the key enzyme that catalyzes the conversion of collagen lysine residuals into allysine, providing the substrate for collagen cross-link formation [74]. Tenascin-C is a glycoprotein that modulates cell-matrix interaction by attenuating the activation of focal adhesion kinase (FAK) and downstream RhoA pathway [75]. These changes signify the reciprocal cellular response to the impaired PCM microenvironment.

In meniscus tissue engineering, mechanical stimulations are often employed to augment the biosynthesis of meniscal cells and quality of engineered products [28]. Building upon this study, harnessing the molecular assembly and mechanobiology of PCM could enhance the responsiveness of meniscal cells to biomechanical stimuli. Notably, in cartilage repair, the application of chondrocytes along with their native PCM, referred to as “chondrons”, has demonstrated positive clinical impacts in commercial applications [76]. Given the pivotal role of collagen V in meniscus PCM highlighted by this study (Fig. 6), the development of collagen V-targeting gene therapy or biomaterials could pave the way for novel strategies to further advance meniscus regeneration. Considering that the PCM is a prevalent feature across various fibrocartilage types, the concept of modulating the PCM or its constituents also holds promise for the regeneration and disease intervention of other fibrocartilaginous tissues, such as the annulus fibrosus [26], disc endplate (Fig. 1c) and TMJ condylar cartilage [77]. In addition, while the *Col5a1^+/−^* meniscus serves as a model of defective PCM, it also represents classic Ehlers-Danlos syndrome (cEDS), a heritable genetic disease with a prevalence of 1:20,000, caused by mutations or haploinsufficiency of the *COL5A1* or *COL5A2* gene [78]. Consequently, our results provide new insights into the higher joint instability and OA propensity observed in cEDS patients [79], potentially guiding improved patient care [80].

## CONCLUSIONS

This study identifies the pericellular matrix (PCM) as a structurally distinct microdomain that envelopes individual cells within fibrocartilage. Utilizing the murine meniscus as a model system, we demonstrate that the PCM is characterized by a random, porous collagen fibrillar network and preferred localization of proteoglycans. This structure confers distinct micromechanical properties compared to the bulk ECM, endowing the PCM with mechanoprotection of resident cells under a complex interplay of tensile and compressive loads. Studying the collagen V-deficient model, we underscore the crucial role of collagen V in mediating the fibrillar structural integrity, mechanical properties and mechanobiological functions of both native PCM and neo-PCM synthesized by meniscal cells in vitro. These findings establish collagen V as a promising target for modulating PCM assembly and cell-matrix interactions in meniscus regeneration and repair. Given the prevalence of PCM in other fibrocartilaginous tissues, the development of new regeneration or disease intervention strategies focusing on the PCM holds considerable promise for clinical translation in addressing various fibrocartilage-associated diseases.

## METHODS

### Human and Bovine Specimens

Human meniscus, articular cartilage, patella tendon and intervertebral disc specimens were acquired from de-identified, healthy donors through National Disease Research Interchange (NDRI, *n* = 3 for each tissue). All human sample-related experiments were exempted for review by the determination of Drexel University Institutional Review Boards (IRB). Bovine medial menisci were harvested from knee joints of adult cows (18-30 months of age, *n* = 3, Animal Technologies). Samples were prepared for histology, immunofluorescence (IF) imaging and collagen nanostructure analysis.

### Murine Model

Wild-type (WT) and collagen V-haploinsufficient (*Col5a1^+/−^* [34]) mice in the C57BL/6 background were housed in the Calhoun animal facility at Drexel University. Animal use and care were approved by the Institutional Animal Care and Use Committee (IACUC) at Drexel University, following the NIH Guide for the Care and Use of Laboratory Animals. Medial menisci from 3-month-old *Col5a1^+/−^* and WT littermates were harvested and prepared for electron microscopy, AFM-based nanoindentation and force relaxation tests, and intracellular calcium signaling assays. In addition, medial menisci from 3-week-old *Col5a1^+/−^* and WT littermates were prepared for intracellular calcium signaling assays, NanoString multiplex gene expression analysis and in vitro cell studies. Both male and female mice were included, as we did not detect significant sex-associated differences in the histological or biomechanical properties of the meniscus.

### Histology and Immunofluorescence Imaging

Safranin-O/Fast Green staining was applied to assess tissue morphology and sulfated glycosaminoglycan (sGAG) distribution, and IF imaging was applied to assess the presence and distribution of specific matrix molecules. Paraffin embedded human samples were sectioned via microtomy at 6 µm-thickness on both transverse and sagittal planes of the meniscus, transverse plane of intervertebral disc, sagittal plane of articular cartilage, and frontal plane of patella tendon. Optimal Cutting Temperature medium (OCT) embedded bovine and murine menisci were cryo-sectioned into 6-µm-thick slices along the sagittal plane for bovine samples, and both transverse and sagittal planes for murine samples via the Kawamoto’s film method [81]. For IF imaging, sections were fixed in 4% paraformaldehyde (PFA) for 10 min, washed in PBS, blocked 30 min in 5% BSA 1% goat serum buffer, followed by incubation of primary antibodies (perlecan: A7L6, 1:200, Santa Cruz Biotech; collagen V: AB7046, 1:200 dilution, Abcam; collagen VI: 70R-CR009X, 1:200, Fitzgerald; aggrecan: AB1031, 1:200, MilliporeSigma; biglycan: LF-159, 1:100, MilliporeSigma) at 4°C overnight. The next day, sections were washed in PBS, incubated with secondary antibodies (perlecan: A-11006, 1:200, Invitrogen; collagen V, collagen VI, aggrecan, biglycan: 65-6120, 1:500, Invitrogen) at room temperature for 2 h, followed by DAPI (0100-20, SouthernBiotech) medium mounting prior to imaging (Leica DMI-6000B). The specificity of collagen V antibody (AB7046) was confirmed via western blot (Fig. S1). Internal negative controls were prepared following the same procedure, except without primary antibody incubation.

### Collagen Fibril Nanostructural Analysis

Scanning electron microscopy (SEM) was performed to visualize the collagen fibril nanostructure on the sections of human meniscus, articular cartilage, tendon, intervertebral disc, bovine meniscus, as well as 3-month-old WT murine meniscus. Six-µm-thick cryo-sections were prepared from the transverse plane of human, bovine and murine meniscus, human intervertebral disc, sagittal plane of human articular cartilage, and frontal plane of human patella tendon. Sections were treated with 0.1% trypsin (T7409, Sigma), followed by 0.1% hyaluronidase (H3506, Sigma) at 37°C for 24 h each, to remove proteoglycans and non-fibrillar constituents. Samples were then fixed with Karnovsky’s fixative (Electron Microscopy Sciences) at room temperature for 3 h, dehydrated in a series of graded water-ethanol and ethanol-hexamethyldisilazane (HMDS, A15139, Alfa Aesar) mixtures, and air dried overnight, following the established procedure [82]. SEM (Zeiss Supra 50VP) images were acquired on samples coated with ≈ 6 nm thick platinum.

Serial block-face scanning electron microscopy (SBF-SEM) was applied to highlight the distinct 3-dimensional fibril architecture of the PCM in 3-month-old WT murine meniscus. Freshly dissected menisci were fixed in Karnovsky’s fixative for 15 min at room temperature, then placed on orbital shaker with gentle movement for 2 h at 4°C. Menisci were kept in fixation solution and shipped to the University of Delaware overnight on ice without being frozen, then embedded in resin [83]. Serial sections were performed on the transverse plane of meniscus and imaged by Apreo VolumeScope (ThermoFisher). Within each 30 × 15 × 10 µm^3^ region of interest (ROI), segmentation of the images was performed by Seg3D, then reconstituted into 3D images using Amira (ThermoFisher).

Transmission electron microscopy (TEM) was applied to quantify the collagen fibril diameter of the murine meniscus PCM and ECM. Freshly dissected menisci from 3-month-old WT and *Col5a1^+/−^*mice (*n* = 5/genotype) were fixed in Karnovsky’s fixative for 15 min at room temperature, then placed on orbital shaker with gentle movement for 2 h at 4°C. Menisci were kept in fixation solution and shipped to the University of South Florida overnight on ice without being frozen. Samples were rinsed with sodium cacodylate buffer and post-fixed for 1 h with 1% osmium tetroxide, dehydrated in an ethanol series followed by 100% propylene oxide, infiltrated and embedded for 3 d in a mixture of Embed 812, nadic methyl anhydride, dodecenylsuccinic anhydride and DMP-30 (EM Sciences) and polymerized overnight at 60°C, following the established procedure [82]. Ultra-thin sections, ≈ 90 nm thickness, of the meniscus at frontal plane were prepared using a Leica ultramicrotome and post-stained with 2% aqueous uranyl acetate and 1% phosphotungstic acid, pH 3.2. Sections were examined and imaged at 80 kV using a JEOL 1400 TEM (JEOL) equipped with a Gatan Orius widefield side mount CCD camera (Gatan). Based on the TEM images, collagen fibril diameter and heterogeneity were quantified via ImageJ by two independent researchers.

### IF-guided AFM Nanomechanical Mapping

To quantify the micromechanical properties of meniscus PCM and T/IT-ECM, freshly dissected 3-month-old murine menisci were embedded in OCT medium to obtain 6-µm-thick, unfixed cryo-sections on either horizontal (longitudinal) or vertical (transverse) plane of the meniscus via Kawamoto’s film method [81]. Each cryo-section was washed in PBS to remove excessive OCT, blocked with 10% goat serum for 30 min, and then, immuno-labelled with perlecan, the biomarker of meniscus PCM [84,85] by incubation with the primary perlecan antibody (A7L6, 1:50) followed by secondary antibody (A-11006, 1:200), for 20 min each at room temperature. This immuno-labelling procedure has been shown not to significantly alter the micromechanics of the matrix [86]. On each section, we identified 2 ROIs in the outer zone of the uncalcified meniscus central body. The size of each ROI is 20 × 20 μm^2^, containing well-defined PCM terrains. Guided by perlecan IF-imaging, within each ROI, AFM-nanomechanical mapping was executed in a 40 × 40 indentation grid within 1× PBS at room temperature using microspherical tips (*R* ≈ 2.25 µm, *k* ≈ 0.6 N/m, µMasch) up to ≈ 60 nN maximum indentation force at 10 μm/s indentation rate using an MFP-3D AFM (Asylum Research), following the established procedure [82,87,88]. The effective indentation modulus, *E*_ind_, was calculated by fitting the entire loading portion of each indentation force-distance (*F-D*) curve to the finite thickness-corrected Hertz model [89], assuming the Poisson’s ratio ν ≈ 0 for the meniscus [90].

Within the same ROI, to quantify the viscoelastic micromechanics, ramp-and-hold force relaxation test was executed in a 6 × 6 grid. Following the same procedure as nanoindentation, the tip was programmed to indent the sample up to ≈ 60 nN maximum force at 10 µm/s z-piezo displacement rate. The *z*-piezo was then held at a constant position for ≈ 30 s, resulting in an approximately constant indentation depth. During this period, both the *z*-piezo displacement and cantilever bending were recorded as a function of time at 500 Hz sampling rate, from which, the temporal profiles of indentation force, *F*(*t*), and depth, *D*(*t*) (mostly constant), were extracted. The temporal modulus, *E*(*t*) was then calculated by fitting *F*(*t*) and *D*(*t*) to the substrate corrected-Hertz model, with the correction for taking into account the finite indentation ramp rate [91,92]. The relaxation of *E*(*t*) was then fitted to the five-element standard linear solid (SLS) model [93],

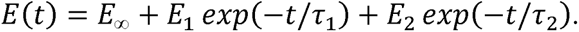

This model yielded the equilibrium modulus, *E*_∞_, and the time-dependent properties corresponding to the two relaxation modes, (*E*_1_, τ_1_) and (*E*_2_, τ_2_) (τ_1_ < τ_2_). The instantaneous indentation modulus, *E*_0_, was estimated as, E_O_ = E_∞_ + E_l_ + E_2_. We used the ratio of equilibrium versus instantaneous modulus, *E*_∞_/*E*_0_, as an indicator of the degree of elasticity. For each outcome, based on the perlecan IF-labelling, we separated the values corresponding to the PCM versus T/IT-ECM using a custom MATLAB program when applicable, and excluded values corresponding to cell remnants (e.g., cellular debris consisting of cytoplasmic organelles and nucleus damaged during sectioning), following the established procedure [38].

### Intracellular [Ca^2+^]*_i_* Signaling Under Osmotic Stimulation

We performed in situ calcium signaling analysis on meniscal cells from WT and *Col5a1^+/−^* mice at 3 weeks and 3 months of age. Freshly dissected menisci were labelled with 15 µM Calbryte^TM^ 520 AM (20650, AAT Bioquest) in DMEM (11995065, Gibco) with 10% (v/v) fetal bovine serum (FBS) and 1% penicillin-streptomycin (10378016, Gibco) for 45 min at 37°C, and then, washed 3 times with DMEM prior to imaging. Intracellular calcium signaling, [Ca^2+^]*_i_*, images of cells in the outer region of the non-ossified central body were captured every 1.5 s for 15 min in hypotonic (165 mOsm), isotonic (310 mOsm) and hypertonic (600 mOsm) DMEMs under a 20× objective lens with a confocal microscope (LSM700, Zeiss) [94]. The pattern of [Ca^2+^]*_i_* oscillation was analyzed based on its fluorescent signal and transient behavior, following the established procedure [94]. In brief, the average fluorescent intensity of each cell in the images was extracted and normalized to the basal intensity value when the cell was at rest. A responsive cell was considered as any cell displaying [Ca^2+^]*_i_* signals with the peak signal exceeding four times the maximum fluctuation of the baseline. The percentage of responding cells, %*R*_cell_, was defined as the proportion of responsive cells relative to the total number of cells within a ROI. The number of [Ca^2+^]*_i_* peaks during the 15-min recording period, *n*_peak_, was extracted from all the responsive cells.

### Gene Expression Analysis via NanoString and qPCR

We assessed the expressions of key genes in the meniscal cells using the NanoString multiplex gene expression analysis. Total RNA was extracted from the central body cells of 3-week-old WT and *Col5a1^+/−^* menisci by homogenizing freshly dissected tissues in TRI-reagent (T9424, Sigma) and phase-separated in 1-bromo-3-chloropropane (B9673, Sigma) [82,95]. A total of eight menisci from two siblings for each genotype were pooled as one biological repeat (*n* = 4 biological repeats/genotype) to obtain sufficient RNA. Prior to the analysis, all RNA samples were assessed to validate the purity with a 260/280 ratio of 2.0 to 2.2 by Infinite 200 PRO (Tecan) and the quality with an RNA integrity number (RIN) > 7.0 by 2100 Bioanalyzer (Agilent). NanoString multiplex gene expression analysis was performed using 100 ng RNA for each biological repeat and custom-designed panels of 103 strategically selected genes (Table S5). These genes include 1) collagens [96], 2) collagens post-translational modification (PTM) [97,98], 3) proteoglycans [85,99], 4) other proteins and glycoproteins [99–103], 5) matrix remodeling [104], 6) cell-matrix interactions (integrins [105], cadherins [106], transmembrane ion channels [107,108], and focal adhesions [109,110]), 7) cell surface markers, including adhesion molecules (*Cd44*, *Cd146*) and receptor molecules (*Cd90*, *Cd105*) [111], 8) transcriptional factors [68,112–114]. In addition, we included the major signaling pathways, such as Wnt/β-Catenin (embryonic development, tissue homeostasis, *Bmp2*, *4*, *Dkk3*) [115,116], Connective tissue growth factor (cell proliferation, angiogenesis, tissue fibrosis, *Ctgf*) [117], RhoA (mesenchymal cell differentiation, joint tissue morphogenesis, *Gdf5*) [118,119], Insulin-like growth factors (mammalian growth, development, aging, *Igf1*, *Igfbp2*, *3*) [120], Parathyroid related protein [121], Fibroblast Growth Factor 3 (growth plate proliferation, chondrocyte differentiation, *Ihh*, *Pthlh*) [121], Transforming growth factor-β (physiological embryogenesis, adult tissue homeostasis, *Tgfb1*, *2*, *3*, *r2*) [122,123], Tumor necrosis factor (induction of apoptotic cell death, cell survival and proliferation, *Tnfsf11*, *11b*) [124,125], and Yap/Taz (mechanotransduction, *Yap*, *Taz*) [126]. We also included the genes of Notch (*Notch1*) and CALCR (*Calcr*), which has been shown to be impacted by the loss of collagen V in muscle satellite stem cells [68]. Outcomes were first normalized to internal positive and negative controls and subsequently normalized to the geometric mean of four housekeeping genes *Abl1*, *Actb*, *Gapdh* and *Rps17* [127–129]. Heatmap was then generated using nSolver 4.0 (Nanostring).

Quantitative RT-PCR (qPCR) was performed to validate changes in the expressions of major genes detected by NanoString, including *Col5a1*, *Tnc* and *Lox* (primer sequences listed in Table S6). For each biological repeat, 100 ng RNA per sample was extracted following the same procedure (*n* = 4 biological repeats/genotype), and subjected to reverse transcription using the TaqMan reverse transcription kit (N8080234, ThermoFisher), with amplification via the PowerUp SYBR Green Master Mix (A25742, ThermoFisher) on a RealPlex 4S master cycler (Eppendorf AG).

### Composition and Micromechanics of the Neo-PCM of Meniscal Cells

We extracted cells from the central body of freshly dissected menisci from 3-week-old WT and *Col5a1^+/−^* mice by 0.2% collagenase (CLS-2, Worthington) digestion for 5 h at 37°C, filtered extraction solution through a 70-µm cell strainer (22363548, Fisher Scientific). The extracted cells were cultured overnight in cell culture flask (690 170, Greiner Bio-One) using DMEM (11995065, Gibco) supplemented with 10% FBS and 1% penicillin-streptomycin, followed by the next-day media change to remove residual collagenase [130]. For each genotype, cells extracted from a total of eight menisci from two sibling mice were pooled as one biological replicate and cultured in one flask.

For IF imaging, followed by monolayer culture (174900, ThermoFisher) for 7 days in the same DMEM with media changed every other day, cells were fixed in 4% PFA for 10 min, washed in 1× PBS for 5 min, blocked with 5% BSA and 1% Goat Serum buffer for 30 min, followed by the incubation of primary antibodies (collagen V: AB7046, 1:200 dilution, Abcam; tenascin-C: AB108930, 1:200, Abcam; LOX: NBP2-24877, 1:100, Novus Biologicals) at 4°C overnight. Samples were washed with PBS, incubated with the secondary antibody (65-6120, 1:500, Invitrogen) at room temperature for 2 h, washed again with PBS, and then mounted with DAPI prior to imaging (Leica DMI-6000B). Internal negative controls were prepared following the same procedure, without the incubation of primary antibodies.

To test the micromechanical properties of meniscal cells and newly synthesized PCM, 35-mm petri dish (351008, Life Sciences) was treated with 20 µg/mL human fibronectin (10838039001, Sigma-Aldrich) for 12 h prior to seeding meniscal cells with an initial density of 5,000 cells per dish. Cells were fed with DMEM (11995065, Gibco) with 10% FBS and 1% penicillin-streptomycin. AFM-nanoindentation was applied to individual meniscal cells at culture day 0 and 7 to quantify the micromechanics of meniscal cells without and with the newly synthesized matrix, respectively. For the test at day 0, cells were seeded for 3 hr prior to testing to allow for cell adhesion and equilibration. The nanoindentation test was performed using microspherical tips (*k* ≈ 0.03 N/m, *R* ≈ 5 µm, HQ:CSC38/tipless/Cr-Au, cantilever B, NanoAndMore) up to ≈ 12 nN maximum indentation force at 10 μm/s indentation rate using a Dimension Icon AFM (Bruker Nano). The effective indentation modulus, *E*_ind_, was calculated by fitting the entire loading portion of each *F-D* curve to the finite thickness-corrected Hertz model, assuming Poisson’s ratios of 0.38 for cell [131] and 0.04 for newly synthesized cellular matrix [132], respectively, and ≈ 5 µm for the height of the cell [133] and ≈ 9 µm for the height of the cell-matrix composite.

### Fabrication of Aligned Electrospun Nanofibrous Scaffolds

To test if deficiency of collagen V impairs the strain transmission of the PCM under applied tensile deformation, meniscal cells were additionally cultured on aligned poly(ε-caprolactone) (PCL) nanofibrous scaffold. First, composite fiber-aligned fibrous scaffolds consisting of PCL and poly (ethylene oxide) (PEO) were produced by dual-component electrospinning as previously described [134,135]. The use of PEO here was to provide a sacrificial fiber constituent to increase the scaffold porosity and allow the infiltration of meniscal cells during culture. In brief, PCL (80 kDa, Sigma Aldrich) was dissolved in a 1:1 mixture of tetrahydrofuran and N,N-dimethylformamide to yield a 14.3% w/v solution (Fisher Chemical), and PEO (200 kDa, Polysciences) was dissolved in 90% ethanol to yield a 10% w/v solution. The polymer solutions were electrospun to generate 250 µm-thick aligned composite fiber sheets of 50:50 PCL and PEO using a custom electrospinning device. These scaffolds were then immersed in DI water to remove the sacrificial PEO fibers, hydrated and sterilized through a gradient of ethanol (100%, 70%, 50%, 30%), then coated with 20 μg/mL fibronectin for 12 h.

### Click Labeling of Nascent Proteins in the Neo-PCM of Meniscal Cells

To image the presence and distribution of newly synthesized proteins in the neo-matrix, following the extraction, meniscal cells were cultured at a seeding density of 5 × 10^3^ on a 1 × 1 cm^2^ PCL nanofibrous scaffold up to 7 days, fed by the “azidohomoalanine (AHA) media” with glutamine-, methionine-, and cystine-free DMEM (21013-024, Gibco), supplemented with 1 × 10^−4^ M AHA (1066-25, Click Chemistry Tools), 4 × 10^−3^ M GlutaMAX supplement (35050061, Gibco), 2 × 10^−4^ M L-cystine (C7602, Sigma-Aldrich), 100 μg/mL sodium pyruvate (11360070, Gibco), 50 µg/mL ascorbic acid-2-phosphate (A8960, Sigma-Aldrich), 10% (v/v) FBS and 1% penicillin-streptomycin [136]. To label newly synthesized matrix, at day 1, 3 and 7 of culture, cells were stained by fluorophore-conjugated cyclooctyne (DBCO-488, 30 µM) in 1% BSA at 37 °C for 30 min, washed with 1× PBS wash, and fixed in 4% PFA for 30 min at room temperature, and stained for plasma membrane (CellMask Deep Red, 1:1000, Invitrogen) prior to imaging. Fluorescence images were taken using a confocal microscope (LSM700, Zeiss) [136]. ImageJ was used to quantify the nascent matrix thickness, masked by the outer edge of cell membrane, the midsection of each cell was measured radially (*n* = 5 measurements per cell) as the distance that the matrix extended, the average of 5 measurements from each cell was used for nascent matrix thickness [136].

### Deformation of Meniscal Cells and Neo-PCM Under Applied Tensile Strain

To test the strain transmission of the neo-PCM under applied tensile strain, the meniscal cell-seeded PCL scaffolds were subjected to tensile deformation using a custom tensile strain control platform integrated with the multiphoton (LSM 880, Zeiss) confocal microscope (Fig. S5). A linear stage with two arms was designed with one arm fixed, and the other mobile arm was driven by a one-axis high load tilt platform (PT-QX03, PDV) with a caliper micrometer head to ensure linear, gradual and accurate movement application at an increment of 20 μm. A 1 × 0.3 cm^2^ PCL scaffold was seeded with meniscal cells, cultured for 7 days in “AHA media”, and then, mounted onto the two arms of the platform, with the central 5 mm region being the active loading domain. Prior to the stretch, a 30 µm pre-stretch was applied to minimize any sample-motor contact slacks. Then, a 1 mm tensile stretch was applied to the cell-seeded PCL scaffold, resulting in ≈ 20% apparent tensile strain in the scaffold. The system was then equilibrated for 30 seconds prior to imaging. Multichannel *z*-stack confocal images were taken on the visually tracked same ROI both immediately before and after the application of tensile stretch. During the application of tensile stretch and imaging, the scaffold was kept in DMEM to maintain the near physiological environment. ImageJ was used to quantify the dimensions of the nascent protein layer and cell membrane within the actively loaded region.

### Statistical Analysis

Linear statistical models were applied to analyze *E*_ind_ (tissue), *E*_0_, *E*_∞_, *E*_1_, *E*_2_, τ_1_, τ_2_, ratio, *d*_col_, *%R*_cell_, *n*_peak_, nascent protein thickness, and *E*_ind_ (cell), using the R package lme4 (version 1.1-27.1) [137]. For continuous dependent variables, *E*_ind_ (tissue), *E*_0_, *E*_∞_, *E*_1_, *E*_2_, τ_1_, τ_2_, ratio, *d*_col_, nascent protein thickness and *E*_ind_ (cell), the linear mixed effect model (LMM) was applied. For non-continuous variables, the generalized linear model (GLMM) was applied to *%R*_cell_ (binary) with the binomial family and *n*_peak_ (count) with the Poisson family, respectively. In these tests, genotypes (*+/+* versus *+/−*), anisotropy (horizontal versus vertical), region (PCM versus ECM), age (3-week-old versus 3-month-old), osmolarity condition (hypo- versus iso- versus hypertonic) were treated as fixed effect factors when applicable, while individual animal/cell effect was treated as a randomized factor. Prior to applying linear models, Shapiro-Wilk test was applied to residuals to confirm that outcomes did not significantly deviate from normal distribution, and likelihood ratio test was applied to determine the covariance structure of the data, unstructured versus compound symmetry. For collagen structural data, F-test was applied to compare fibril diameter variances. DESeq2 method was applied to test the gene express difference the genotypes. Unpaired two-sample student’s *t*-test was applied to examine the differences in qPCR, strain transmission outcomes. Prior to applying student’s *t*-test, Shapiro-Wilk test was applied to confirm that outcomes follow normal distribution. The significance level was set at α = 0.05. All quantitative and statistical outcomes were summarized in Tables S1-S4, S6 and S7.

## ACKNOWLEDGEMENTS

This work was financially supported by the National Science Foundation (NSF) Grant CMMI-1751898 (to LH), National Institutes of Health (NIH) Grant R01AR074490 (to LH), R01AR075418 (to RLM and LH), R01AR056624 (to JAB and RLM) and R01AR074472 (to XLL), as well as NIH Grant P30AR069619 to the Penn Center for Musculoskeletal Disorders. We thank the Singh Center for Nanotechnology at the University of Pennsylvania for the use of TIRF MFP-3D, which is part of the National Nanotechnology Coordinated Infrastructure Program supported by the NSF Grant NNCI-1542153. We thank Dr. Vivek B. Shenoy (University of Pennsylvania) for valuable discussions, and Dr. Russell Fernandes (University of Washington) for the kind gift of collagen XI extracted from fetal bovine cartilage. All imaging was performed at Drexel University’s Cell Imaging Center.

