## Supporting Materials for "Structure-Mechanics Principles and Mechanobiology of Fibrocartilage Pericellular Matrix: A Pivotal Role of Type V Collagen"

*Correspondence and requests for materials should be addressed to:

Dr. Lin Han

**Table S1.** Summary of averaged values and statistical analysis outcomes of the micromodulus, $E_{ind}$, between pericellular matrix (PCM) and extracellular matrix (ECM) for 3-month-old wild-type (WT) meniscus at each region (PCM versus ECM) and anisotropy (horizontal versus vertical), shown as mean ± 95% confidence interval (CI) from values averaged by each animal.

| | PCM | | | | *p*-value | ECM | | | | *p*-value |
|---|---|---|---|---|---|---|---|---|---|---|
| | Horizontal | | Vertical | | | Horizontal | | Vertical | | |
| | mean ± 95% CI | *n* | mean ± 95% CI | *n* | | mean ± 95% CI | *n* | mean ± 95% CI | *n* | |
| $E_{ind}$ (kPa) | 247 ± 31 | 9 | 270 ± 71 | 7 | 0.003 | 390 ± 62 | 9 | 538 ± 138 | 7 | < 0.001 |
| $E_0$ (kPa) | 179 ± 33 | 5 | 204 ± 38 | 5 | 0.027 | 298 ± 75 | 5 | 458 ± 170 | 5 | 0.002 |
| $E_\infty$ (kPa) | 128 ± 44 | 5 | 143 ± 27 | 5 | 0.040 | 228 ± 61 | 5 | 373 ± 166 | 5 | < 0.001 |
| $E_1$ (kPa) | 29 ± 17 | 5 | 32 ± 21 | 5 | 0.166 | 43 ± 15 | 5 | 36 ± 24 | 5 | 0.682 |
| $E_2$ (kPa) | 22 ± 12 | 5 | 29 ± 8 | 5 | 0.511 | 26 ± 10 | 5 | 48 ± 16 | 5 | 0.023 |
| $E_\infty/E_0$ | 0.70 ± 0.18 | 5 | 0.71 ± 0.07 | 5 | 0.450 | 0.74 ± 0.08 | 5 | 0.80 ± 0.08 | 5 | 0.209 |
| $\tau_1$ (sec) | 0.50 ± 0.10 | 5 | 0.54 ± 0.16 | 5 | 0.751 | 0.48 ± 0.13 | 5 | 0.67 ± 0.08 | 5 | 0.063 |
| $\tau_2$ (sec) | 22 ± 3 | 5 | 20 ± 3 | 5 | 0.840 | 23 ± 5 | 5 | 20 ± 2 | 5 | 0.671 |

| *p*-value | Horizontal vs. Vertical | |
|---|---|---|
| | PCM | ECM |
| $E_{ind}$ (kPa) | 0.616 | 0.004 |
| $E_0$ (kPa) | 0.612 | 0.006 |
| $E_\infty$ (kPa) | 0.733 | 0.003 |
| $E_1$ (kPa) | 0.776 | 0.577 |
| $E_2$ (kPa) | 0.281 | 0.012 |
| $E_\infty/E_0$ | 0.801 | 0.397 |
| $\tau_1$ (sec) | 0.577 | 0.014 |
| $\tau_2$ (sec) | 0.251 | 0.389 |

**Table S2.** Summary of collagen fibril diameter distributions and statistical analysis outcomes of the meniscus from 3-month-old WT and *Col5a1*$^{+/-}$ mice, as measured by TEM.

| | PCM | | ECM | |
|---|---|---|---|---|
| | WT | *Col5a1*$^{+/-}$ | WT | *Col5a1*$^{+/-}$ |
| mean | 67 | 81 | 94 | 92 |
| std | 30 | 40 | 32 | 37 |
| $Q_1$ | 43 | 47 | 74 | 63 |
| $Q_2$ | 66 | 77 | 98 | 88 |
| $Q_3$ | 89 | 113 | 116 | 118 |
| min | 8 | 8 | 11 | 22 |
| max | 168 | 195 | 183 | 197 |
| $n_{fibrils}$ | 1,369 | 945 | 994 | 644 |

| | mean($d_{col}$) | | var($d_{col}$) | |
|---|---|---|---|---|
| *p*-value (WT vs *Col5a1*$^{+/-}$) | PCM | ECM | PCM | ECM |
| | 0.001 | 0.781 | < 0.001 | < 0.001 |
| *p*-value (PCM vs ECM) | WT | *Col5a1*$^{+/-}$ | WT | *Col5a1*$^{+/-}$ |
| | < 0.001 | < 0.001 | 0.006 | 0.038 |

**Table S3.** Summary of averaged values and statistical analysis outcomes of the micromodulus, $E_{ind}$, between genotype (WT versus *Col5a1*$^{+/-}$) for 3-month-old meniscus at each anisotropy (horizontal versus vertical) and region (PCM versus ECM), shown as mean ± 95% CI from the values averaged by each animal.

| | PCM | | | | | | | | | |
|---|---|---|---|---|---|---|---|---|---|---|
| | Horizontal | | | | *p*-value | Vertical | | | | *p*-value |
| | WT | | *Col5a1*$^{+/-}$ | | | WT | | *Col5a1*$^{+/-}$ | | |
| | mean ± 95% CI | *n* | mean ± 95% CI | *n* | | mean ± 95% CI | *n* | mean ± 95% CI | *n* | |
| $E_{ind}$ (kPa) | 247 ± 31 | 9 | 107 ± 44 | 9 | 0.001 | 270 ± 71 | 7 | 205 ± 57 | 5 | 0.163 |
| $E_0$ (kPa) | 179 ± 33 | 5 | 57 ± 25 | 5 | 0.003 | 204 ± 38 | 5 | 136 ± 28 | 5 | 0.074 |
| $E_\infty$ (kPa) | 128 ± 44 | 5 | 35 ± 17 | 5 | 0.011 | 143 ± 27 | 5 | 93 ± 29 | 5 | 0.156 |
| $E_1$ (kPa) | 29 ± 17 | 5 | 12 ± 9 | 5 | 0.09 | 32 ± 21 | 5 | 15 ± 13 | 5 | 0.14 |
| $E_2$ (kPa) | 22 ± 12 | 5 | 10 ± 7 | 5 | 0.083 | 29 ± 8 | 5 | 25 ± 11 | 5 | 0.535 |
| $E_\infty/E_0$ | 0.70 ± 0.18 | 5 | 0.64 ± 0.12 | 5 | 0.248 | 0.71 ± 0.07 | 5 | 0.70 ± 0.10 | 5 | 0.813 |
| $\tau_1$ (sec) | 0.50 ± 0.10 | 5 | 0.46 ± 0.12 | 5 | 0.594 | 0.54 ± 0.16 | 5 | 0.65 ± 0.23 | 5 | 0.134 |
| $\tau_2$ (sec) | 22 ± 3 | 5 | 15 ± 3 | 5 | < 0.001 | 20 ± 3 | 5 | 15 ± 3 | 5 | 0.007 |

| | ECM | | | | | | | | | |
|---|---|---|---|---|---|---|---|---|---|---|
| | Horizontal | | | | *p*-value | Vertical | | | | *p*-value |
| | WT | | *Col5a1*$^{+/-}$ | | | WT | | *Col5a1*$^{+/-}$ | | |
| | mean ± 95% CI | *n* | mean ± 95% CI | *n* | | mean ± 95% CI | *n* | mean ± 95% CI | *n* | |
| $E_{ind}$ (kPa) | 390 ± 62 | 9 | 157 ± 57 | 9 | < 0.001 | 538 ± 138 | 7 | 312 ± 50 | 5 | < 0.001 |
| $E_0$ (kPa) | 298 ± 75 | 5 | 77 ± 40 | 5 | < 0.001 | 458 ± 170 | 5 | 273 ± 46 | 5 | < 0.001 |
| $E_\infty$ (kPa) | 228 ± 61 | 5 | 48 ± 22 | 5 | < 0.001 | 373 ± 166 | 5 | 198 ± 28 | 5 | < 0.001 |
| $E_1$ (kPa) | 43 ± 15 | 5 | 17 ± 6 | 5 | 0.006 | 36 ± 24 | 5 | 31 ± 30 | 5 | 0.544 |
| $E_2$ (kPa) | 26 ± 10 | 5 | 14 ± 11 | 5 | 0.074 | 48 ± 16 | 5 | 44 ± 25 | 5 | 0.549 |
| $E_\infty/E_0$ | 0.74 ± 0.08 | 5 | 0.66 ± 0.11 | 5 | 0.109 | 0.80 ± 0.08 | 5 | 0.73 ± 0.12 | 5 | 0.192 |
| $\tau_1$ (sec) | 0.48 ± 0.13 | 5 | 0.42 ± 0.16 | 5 | 0.436 | 0.67 ± 0.08 | 5 | 0.71 ± 0.14 | 5 | 0.654 |
| $\tau_2$ (sec) | 23 ± 5 | 5 | 16 ± 3 | 5 | 0.001 | 20 ± 2 | 5 | 16 ± 4 | 5 | 0.024 |

| *p*-value | Anisotropy (Horizontal vs Vertical) | | | | Region (PCM vs ECM) | | | |
|---|---|---|---|---|---|---|---|---|
| | WT | | *Col5a1*$^{+/-}$ | | WT | | *Col5a1*$^{+/-}$ | |
| | PCM | ECM | PCM | ECM | Horizontal | Vertical | Horizontal | Vertical |
| $E_{ind}$ (kPa) | 0.557 | 0.001 | 0.031 | 0.001 | < 0.001 | < 0.001 | 0.081 | 0.008 |
| $E_0$ (kPa) | 0.500 | < 0.001 | 0.042 | < 0.001 | 0.004 | < 0.001 | 0.588 | 0.001 |
| $E_\infty$ (kPa) | 0.653 | < 0.001 | 0.098 | < 0.001 | 0.007 | < 0.001 | 0.695 | 0.005 |
| $E_1$ (kPa) | 0.760 | 0.480 | 0.590 | 0.110 | 0.074 | 0.526 | 0.696 | 0.096 |
| $E_2$ (kPa) | 0.336 | 0.004 | 0.040 | < 0.001 | 0.476 | 0.004 | 0.520 | 0.004 |
| $E_\infty/E_0$ | 0.779 | 0.270 | 0.231 | 0.160 | 0.188 | 0.014 | 0.552 | 0.352 |
| $\tau_1$ (sec) | 0.636 | 0.016 | 0.016 | 0.001 | 0.783 | 0.083 | 0.597 | 0.477 |
| $\tau_2$ (sec) | 0.201 | 0.208 | 0.871 | 0.905 | 0.686 | 0.66 | 0.143 | 0.163 |

**Table S4.** Summary of averaged values and statistical analysis outcomes of $[Ca^{2+}]_i$ parameters between genotype (WT versus *Col5a1*$^{+/-}$) for meniscal cells at each osmolarity (hypotonic versus isotonic versus hypertonic) and age (3-week versus 3-month), shown as mean ± 95% CI from the values averaged by cells.

| | $\%R_{cell}$ | | | | | |
|---|---|---|---|---|---|---|
| | 3 weeks | | | 3 months | | |
| mean ± 95% CI | Hypotonic | Isotonic | Hypertonic | Hypotonic | Isotonic | Hypertonic |
| WT | 74 ± 5 | 60 ± 6 | 26 ± 5 | 63 ± 5 | 43 ± 5 | 22 ± 4 |
| *Col5a1*$^{+/-}$ | 50 ± 5 | 45 ± 5 | 22 ± 4 | 45 ± 5 | 35 ± 5 | 15 ± 3 |
| *p*-value (genotype) | < 0.001 | < 0.001 | 0.265 | < 0.001 | 0.024 | 0.009 |
| *p*-value (osmolarity) | Hypo vs Iso | Hypo vs Hyper | Iso vs Hyper | Hypo vs Iso | Hypo vs Hyper | Iso vs Hyper |
| WT | < 0.001 | < 0.001 | < 0.001 | < 0.001 | < 0.001 | < 0.001 |
| *Col5a1*$^{+/-}$ | 0.391 | < 0.001 | < 0.001 | 0.023 | < 0.001 | < 0.001 |

| | $n_{peak}$ | | | | | |
|---|---|---|---|---|---|---|
| | 3 weeks | | | 3 months | | |
| mean ± 95% CI | Hypotonic | Isotonic | Hypertonic | Hypotonic | Isotonic | Hypertonic |
| WT | 3.96 ± 0.20 | 4.35 ± 0.39 | 1.09 ± 0.07 | 2.44 ± 0.17 | 1.83 ± 0.16 | 1.07 ± 0.07 |
| *Col5a1*$^{+/-}$ | 3.83 ± 0.26 | 2.32 ± 0.19 | 1.25 ± 0.16 | 2.05 ± 0.19 | 1.61 ± 0.16 | 1.12 ± 0.09 |
| *p*-value (genotype) | 0.487 | < 0.001 | 0.358 | 0.017 | 0.166 | 0.788 |
| *p*-value (osmolarity) | Hypo vs Iso | Hypo vs Hyper | Iso vs Hyper | Hypo vs Iso | Hypo vs Hyper | Iso vs Hyper |
| WT | 0.122 | < 0.001 | < 0.001 | < 0.001 | < 0.001 | < 0.001 |
| *Col5a1*$^{+/-}$ | < 0.001 | < 0.001 | < 0.001 | 0.028 | < 0.001 | 0.023 |

| *p*-value (age) | $\%R_{cell}$ | | | $n_{peak}$ | | |
|---|---|---|---|---|---|---|
| | Hypo | Iso | Hyper | Hypo | Iso | Hyper |
| WT | 0.001 | < 0.001 | 0.251 | < 0.001 | < 0.001 | 0.902 |
| *Col5a1*$^{+/-}$ | 0.188 | 0.006 | 0.008 | < 0.001 | < 0.001 | 0.479 |

**Table S5.** List of genes for multiplex gene expression analysis via Nanostring.

| Collagens | Collagen PTM | Proteoglycans | Other proteins/ glycoproteins | Matrix remodeling |
|---|---|---|---|---|
| *Col1a1, a2* | *Plod1,2a,3* | *Acan* | *Acta2* | *Adamts1,4,5* |
| *Col2a1* | *Lox* | *Bgn* | *Alpl* | *MMP2,3,9,13* |
| *Col3a1* | *Loxl2* | *Dcn* | *Comp* | *Timp1,3* |
| *Col5a1,a2,a3* | *Adamts2* | *Fmod* | *Dmp1* | |
| *Col6a1,a2,a3* | *Bmp1* | *Hspg2* | *Fn1* | |
| *Col9a1,a2,a3* | | *Lum* | *Halpn1* | |
| *Col10a1* | | *Vcan* | *Has2* | |
| *Col11a1,a2* | | | *Matn1,3* | |
| | | | *Prg4* | |
| | | | *Tnc* | |
| **Cell-matrix interactions** | **Cell surface markers** | **Transcription factors** | **Signaling factors** | **Housekeeping** |
| *Cdh2,3,5,11* | *Cd44* | *Mki67* | *Bmp2,4* | *Abl1* |
| *Itga1,3,5,v* | *Cd90* | *Mkx* | *Calcr* | *Actb* |
| *Itgb1,3,5* | *Cd105* | *Pcna* | *Ctgf* | *Gapdh* |
| *Piezo1,2* | *Cd146* | *Pparg* | *Dkk3* | *Rps17* |
| *Trpv4* | | *Rbpj* | *Gdf5* | |
| *Postn* | | *Runx2* | *Igf1* | |
| *Pxn* | | *Scx* | *Igfbp2,3* | |
| *Vcl* | | *Sox9* | *Ihh* | |
| | | | *Notch1* | |
| | | | *Pthlh* | |
| | | | *Tgfb1,2,3,r2* | |
| | | | *Tnfsf11,11b* | |
| | | | *Yap1, Taz* | |

**Table S6.** List of primers for qPCR and a summary of averaged values and statistical analysis outcomes of qPCR between genotype (WT versus *Col5a1*$^{+/-}$) for 3-week-old meniscus.

| Gene | Forward Primer | Reverse Primer |
|---|---|---|
| *Col5a1* | 5'-AAGCGTGGGAAACTGCTCTCCTAT-3' | 5'-AGCAGTTGTAGGTGACGTTCTGGT-3' |
| *Lox* | 5'-ACGGCTACCACAGAAGCTG-3' | 5'-ATGGCTGTTGTTGCTATGGCA-3' |
| *Tnc* | 5'-TCTTCTGCTGCGTGACAACC-3' | 5'-GAGAAACCAGCTTGGAACCAG-3' |
| *β-actin* | 5'-AGATGACCCAGATCATGTTTGAGA-3' | 5'-CACAGCCTGGATGGCTACGT-3' |

| | WT | | *Col5a1*$^{+/-}$ | | *p*-value (genotype) |
|---|---|---|---|---|---|
| | Mean ± 95% CI | *n* | Mean ± 95% CI | *n* | |
| *Col5a1* | 1.00 ± 0.30 | 4 | 0.58 ± 0.30 | 4 | 0.020 |
| *Lox* | 1.00 ± 0.82 | 4 | 2.08 ± 0.74 | 4 | 0.021 |
| *Tnc* | 1.00 ± 0.46 | 4 | 2.24 ± 1.14 | 4 | 0.034 |

**Table S7.** Summary and statistical analysis outcomes of in-vitro analysis between genotypes (WT versus *Col5a1*$^{+/-}$) for 3-week-old meniscus.

| | | WT | | *Col5a1*$^{+/-}$ | | $p$-value (genotype) |
|---|---|---|---|---|---|---|
| | | mean ± 95% CI | $n_{cell}$ | mean ± 95% CI | $n_{cell}$ | |
| Nascent protein thickness (μm) | day 1 | 0.90 ± 0.21 | 31 | 1.02 ± 0.12 | 32 | 0.644 |
| | day 3 | 3.37 ± 0.18 | 40 | 3.91 ± 0.34 | 46 | 0.011 |
| | day 7 | 4.37 ± 0.44 | 32 | 4.14 ± 0.43 | 36 | 0.381 |
| neo-PCM modulus (kPa) | day 0 | 1.34 ± 0.36 | 23 | 1.03 ± 0.37 | 24 | 0.663 |
| | day 7 | 6.25 ± 1.36 | 34 | 1.83 ± 0.57 | 35 | < 0.001 |
| Tensile strain | Nascent protein | 10.10 ± 0.82 | 40 | 12.83 ± 0.68 | 38 | < 0.001 |
| | Cell membrane | 3.67 ± 0.51 | 40 | 4.77 ± 0.63 | 38 | 0.008 |

| $p$-value (time points) | Nascent protein thickness | | | neo-PCM modulus |
|---|---|---|---|---|
| | day 1 vs 3 | day 1 vs 7 | day 3 vs 7 | day 0 vs 7 |
| WT | < 0.001 | < 0.001 | < 0.001 | < 0.001 |
| *Col5a1*$^{+/-}$ | < 0.001 | < 0.001 | 0.629 | 0.209 |

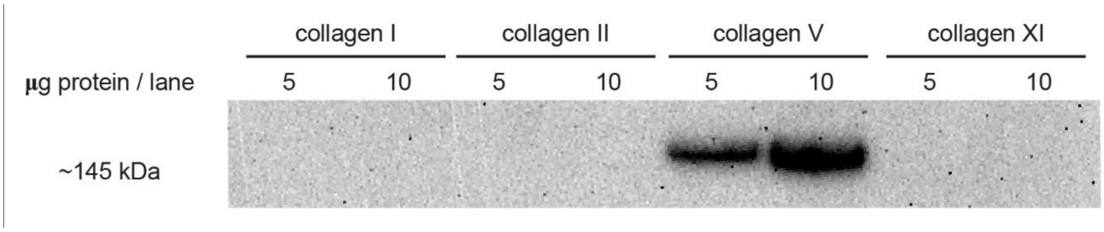


**Figure S1.** Western blot on recombinant mouse collagen I, collagen II extracted from human cartilage, collagen V extracted from bovine skin, collagen XI extracted from fetal bovine cartilage, validated the specificity of collagen V antibody (AB7046).

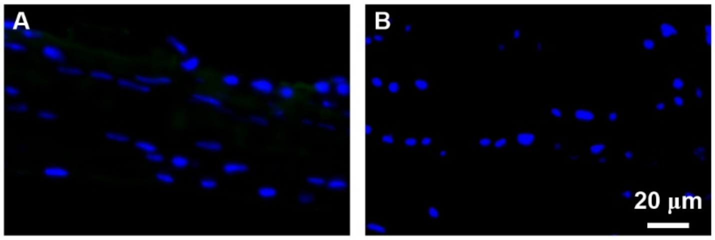


**Figure S2.** Internal negative control for the immunofluorescence imaging analysis. The controls were incubated with secondary antibody but not primary antibody incubation. Panel A: secondary antibody host goat anti-rabbit, B: secondary antibody host goat anti-rat, blue: DAPI.

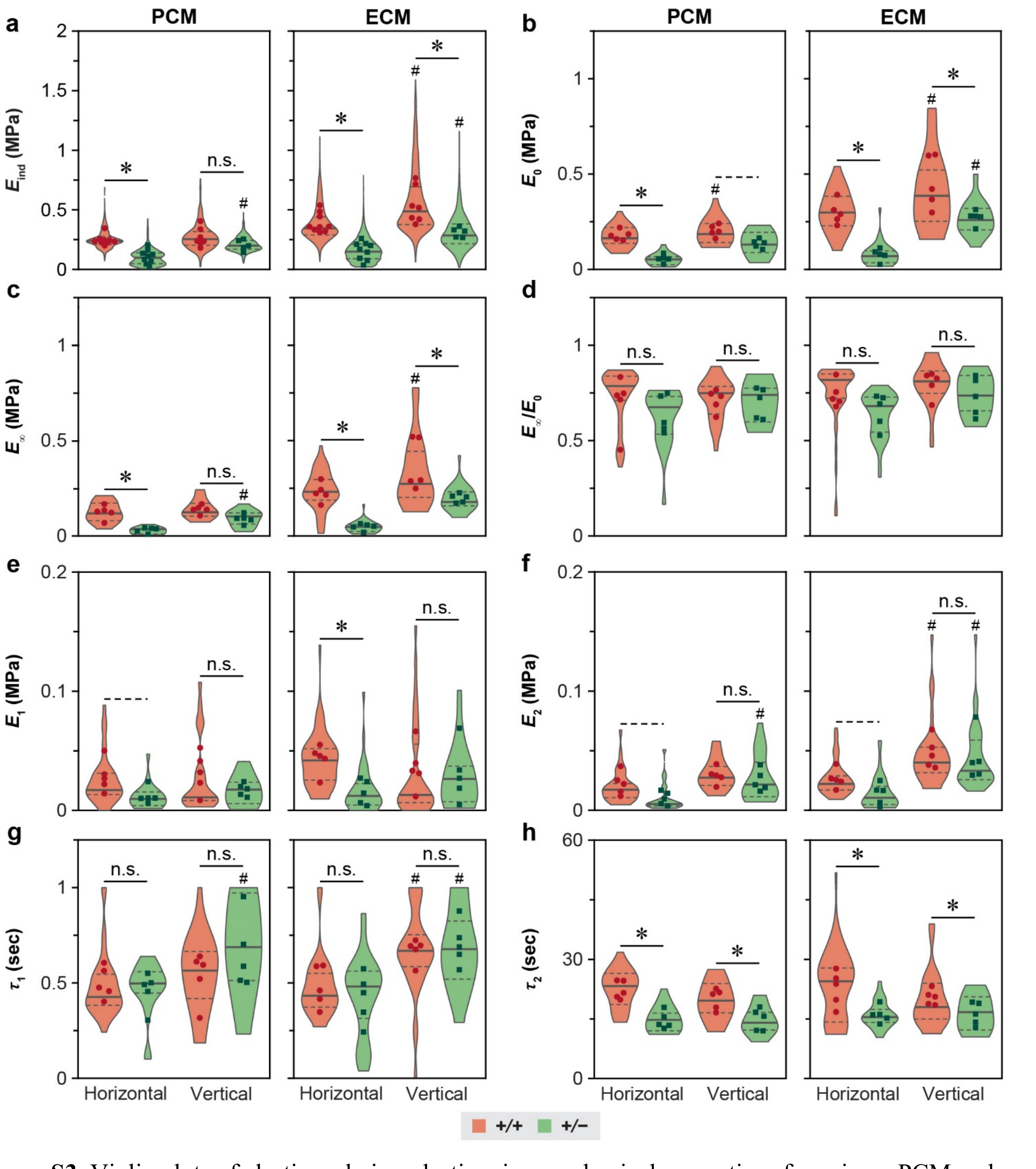


**Figure S3.** Violin plots of elastic and viscoelastic micromechanical properties of meniscus PCM and ECM from 3-month-old WT and *Col5a1*$^{+/-}$ mice on horizontal and vertical cryo-sections, a) indentation micromodulus, $E_{ind}$, b) instantaneous modulus, $E_0$, c) equilibrium modulus, $E_\infty$, d) elasticity ratio, $E_\infty/E_0$, e) short-term viscoelastic relaxation modulus, $E_1$, f) long-term viscoelastic relaxation modulus, $E_2$, g) short-term relaxation time constant, $\tau_1$, and h) long-term relaxation time constant, $\tau_2$. Data are pooled from ≥ 580 positions tested for $E_{ind}$, ≥ 25 positions for viscoelastic properties for each group ($n \geq 5$ mice). Each data point represents the average value measured from one animal. *: $p < 0.05$, dashed line: $p < 0.10$, n.s.: not significant between WT and *Col5a1*$^{+/-}$ groups under same orientation and region, $^{\#}$: $p < 0.05$ between the horizontal and vertical sections for same genotype and region. Results from panels d-h were measured from murine menisci at 3 months of age.

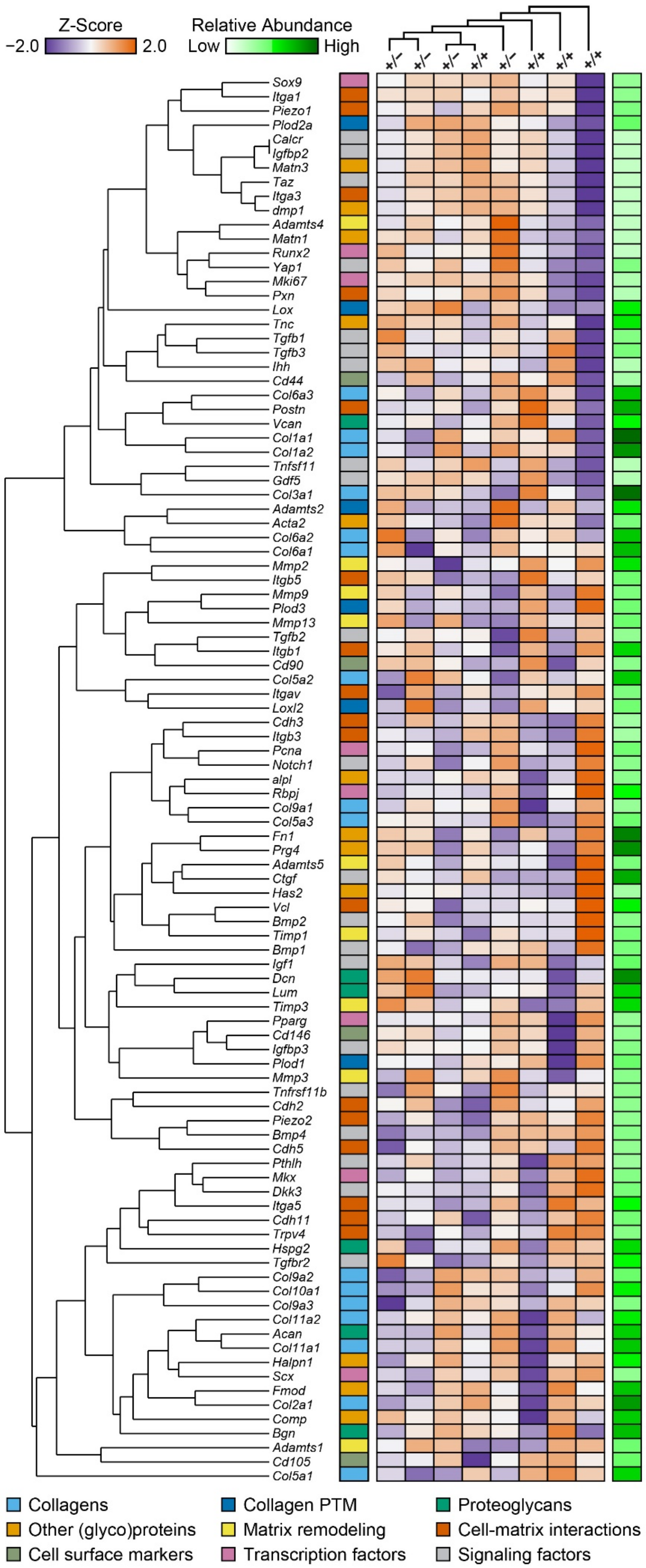


**Figure S4.** Heatmap of clustering of multiplexed full geneset for WT and *Col5a1*$^{+/-}$ mice menisci at 3 weeks of age ($n$ = 4 for each genotype). Color legend represents relative mRNA expression.

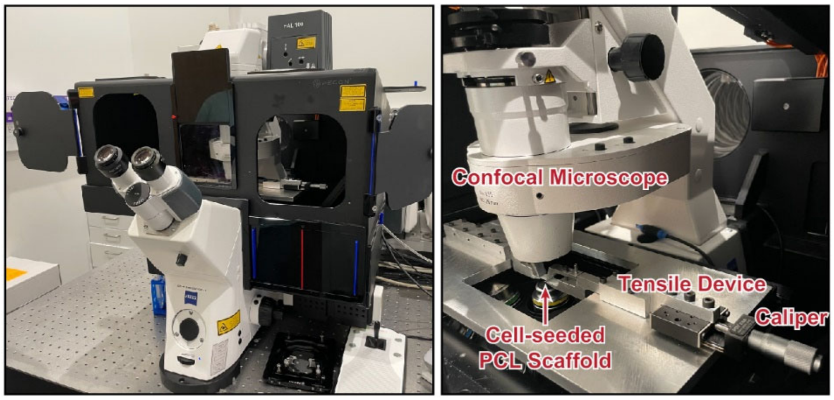


**Figure S5.** Experimental setup of the customized tensile device under the multiphoton confocal imaging system. Cell-seeded PCL was covered by DMEM, and tensile test was controlled by a caliper of the tensile device. The same regions of interest (ROIs) on the PCL before and after 20% tensile strain were imaged by confocal microscope.
